# Sex-biased infections scale to population impacts for an emerging wildlife disease

**DOI:** 10.1101/2022.07.29.502066

**Authors:** Macy J. Kailing, Joseph R. Hoyt, J. Paul White, Heather M. Kaarakka, Jennifer A. Redell, Ariel E. Leon, Tonie E. Rocke, John E. DePue, William H. Scullon, Katy L. Parise, Jeffrey T. Foster, A. Marm Kilpatrick, Kate E. Langwig

## Abstract

Demographic factors are fundamental in shaping infectious disease dynamics. Aspects of populations that create structure, like age and sex, can affect patterns of transmission, infection intensity and population outcomes. However, studies rarely link these processes from individual to population-scale effects. Moreover, the mechanisms underlying demographic differences in disease are frequently unclear. Here, we explore sex-biased infections for a multi-host fungal disease of bats, white-nose syndrome, and link disease-associated mortality between sexes, the distortion of sex ratios, and the potential mechanisms underlying sex differences in infection. We collected data on host traits, infection intensity, and survival of five bat species at 42 sites across seven years. We found females were more infected than males for all five species. Females also had lower apparent survival over winter and accounted for a smaller proportion of populations over time. Notably, female-biased infections were evident by early hibernation and likely driven by sex-based differences in autumn mating behavior. Male bats were more active during autumn which likely reduced replication of the cool-growing fungus. Higher disease impacts in female bats may have cascading effects on bat populations beyond the hibernation season by limiting recruitment and increasing the risk of Allee effects.

## 1. INTRODUCTION

Emerging infectious diseases are a serious threat to wildlife health (1, 2). Population structure can shape epidemics by influencing spatial spread, outbreak size, and host impacts (2–4). Elements of populations that create structure such as classes of individuals of specific ages, sexes, or breeding stages, can have profound effects on disease dynamics (3, 5–9). Sex is an especially important factor because sex-biases in infections can contribute to differential transmission through populations due to behavior (10–15), amplify outbreaks due to seasonal changes in susceptibility (16–19), and modify population impacts through disproportionate mortality (20–22). Differences in infection and mortality can also modulate virulence evolution through sex-specific immune responses that affect pathogen replication and growth (23, 24). As such, determining patterns and mechanisms of sex-biases in infections will improve efforts to minimize outbreaks and manage impacts.

Behavioral and physiological traits are two mechanisms that can produce sex-biases in transmission, susceptibility, infection intensity, or disease-induced mortality (25–27). In most systems, males have an elevated risk of disease compared to females due to territory defense and promiscuous mating, which increases contacts with conspecifics or increase pathogen susceptibility due to physiological stress (9, 25, 28). Sex hormones can also have strong effects on host immunity (29), such that testosterone suppresses immune responses while estrogens enhance it, resulting in weaker immune responses in males, which can increase susceptibility to pathogen infection (25, 30, 31). As such, the majority of empirical studies find that infections are typically male-biased (28, 32, 33), although this generalization is sometimes reversed (34–40) or weak (41) and may be linked to seasonal reproductive stress associated with pregnancy, parturition, or parental care (16, 17). Thus, it is likely that the effects of sex-biased infections may be highly pronounced in disease systems where host reproductive strategies differ seasonally among sexes (19).

White-nose syndrome (WNS) is a highly seasonal fungal disease of bats that has caused population collapse, with declines exceeding 95% in many populations of multiple species (42–46). WNS is caused by the fungal pathogen, *Pseudogymnoascus destructans*, which invades the epidermal tissue of bats during hibernation, and disrupts bat homeostasis, causing water and electrolyte imbalances that increase arousals and deplete stored fat (47–54). Transmission of *P. destructans* occurs through host-to-host contacts and contact between hosts and contaminated environmental reservoirs inside winter hibernation sites (55–59). *Pseudogymnoascus destructans* has a thermal growth range of 0-20°C and optimal growth between 12-16 °C (60). This limits on host growth to the periods when bats are in torpor and they lower their body temperature to the ambient temperatures of their hibernation sites (60–62). The restricted growth of *P. destructans* above 20°C drives the seasonal patterns of WNS, with infection only occurring during winter hibernation and mortality peaking during late winter (56, 59, 63).

Temperate bat species have seasonal sex-biased differences in behaviors that may influence their exposure, susceptibility, and mortality from WNS (64–69). Bats mate in autumn swarms, and females store sperm over winter, delaying ovulation, and giving birth at maternity colonies in spring (70). Male reproductive energy expenditures are highest during autumn when they aggregate at hibernacula (subterranean sites where bats spend the winter) and mate indiscriminately with females (71). Since autumn swarm coincides with the seasonal transmission of *P. destructans* (63), breeding stress among males as well as exposure to the environmental reservoirs (56, 58) could increase their susceptibility to infection. Female energy expenditures are greatest during pregnancy and lactation in spring and summer when hosts typically clear infection (72, 73) and transmission of *P. destructans* is low (63), suggesting females may be the less infected sex. However, female bats may spend more time torpid during winter than males to conserve energy for spring reproduction (66) which could influence the growth of *P. destructans* (60). Due to the differences in reproductive investment and seasonal energy expenditure, sex-biased traits have the potential to affect exposure (e.g., contacting *P. destructans* from other hosts or in the environment) and pathogen growth (e.g., time spent at torpid temperatures that permit fungal growth). Thus, we predict sex-specific behaviors, and the highly seasonal pattern of WNS may drive intersex differences in infection, mortality, and population impacts. Here, we examined differences in *P. destructans* infection between females and males across five bat species in 42 sites across seven years. We then assessed the consequences of intersex differences in infection at both the individual and population-level. Lastly, we explored autumn activity patterns between sexes as a potential mechanism of sex-biased infections.

## 2. MATERIALS AND METHODS

### a. Study sites and sampling design

We sampled bats at hibernacula in Illinois, Massachusetts, Michigan, New York, Vermont, Virginia, and Wisconsin between 2011 and 2021 twice during a seasonal epizootic. Generally, bats were sampled at two time points: early winter (between November and December) and again in late winter (between March and April). During site visits, we walked each section of the hibernacula and counted all the bats of each species and resighted any bats that were previously banded. We sampled up to 20 bats of each species during each visit when possible and stratified sample collection throughout hibernacula to obtain a sample reflective of the spatial distribution of bats at each site. Bat species included little brown (*Myotis lucifugus*), northern long-eared (*Myotis septentrionalis*), tricolored (*Perimyotis subflavus*), big brown (*Eptesicus fuscus*), and Indiana (*Myotis sodalis*) bats. The animal sampling protocols were approved by the Virginia Tech IACUC protocol 17-180. We followed the field decontamination procedures outlined by the United States Fish and Wildlife Service Decontamination Guidelines as well as the recommendations provided by individual states.

### b. *P. destructans* sample collection and quantification

For every bat we sampled, we determined species and biological sex based on external morphology and attached an aluminum-lipped band (2.4, 2.9, and 4.2mm; Porzana Ltd., Icklesham, E. Sussex, U.K.) to the wing for individual identification and resighting. We collected an epidermal swab from individuals to quantify infection prevalence (the fraction of individuals positive for *P. destructans*) and severity (the quantity of *P. destructans* on infected bats) using a standardized swabbing technique (45, 63). We placed swabs in RNAlater® for storage before testing. We extracted DNA and tested for *P. destructans* presence and quantity by qPCR using protocols developed specifically for this fungal pathogen that included a standard growth curve (63, 74).

### c. Activity data collection

At three sites in Wisconsin in 2020, we installed radio frequency identification (RFID) systems, consisting of a passive antenna and an automated data logger (IS10001; Biomark, Boise, ID) at the entrances of each hibernaculum (2-4 per site). The systems at each entrance were equipped with a solar or direct power source to run continuously. During autumn swarm, we captured little brown bats swarming near hibernacula and injected 12.5mm PIT tags (Biomark APT12; Biomark, Boise, ID). Each RFID system was programmed to record each time a unique individual passed through an entrance with a one-minute delay to avoid duplicate detections of tags. We scored a detection as an active bat and used it to characterize activity of females and males.

### d. Statistical analyses

#### i. Infection

We used generalized (GLMM) and linear (LMM) mixed effect models (75) to compare differences in *P. destructans* prevalence and infection severity between males and females of each of the five species. We defined prevalence as the fraction of bats testing positive for *P. destructans* on qPCR (0|1), and infection severity as the log_10_-transformed quantity of fungal DNA (ng) on positive bats (log_10_ fungal loads) which is a well-established metric of infection severity (43, 76, 77). We estimated differences in prevalence between sexes using a GLMM with a binomial distribution and sex and species as fixed effects (Model 1, Appendix 1.1) and compared additive and interactive versions of the model using Akaike Information Criterion (AIC; Appendix 1.2). For prevalence models, we only used data from early hibernation as every bat was infected at the end of hibernation.

We used two datasets to examine changes in fungal loads over winter (between early and late hibernation) between sexes. First, to estimate population-level differences in fungal loads, we used a LMM with fixed additive effects of species and sex and date as a fixed effect interacting with each term (Model 2, Appendix 2.1). Fungal loads are well-correlated with mortality (43, 76, 78), and bats that begin hibernation with higher fungal loads are more likely to die of their infections before the end of winter (79), potentially resulting in survival biases in population-level analyses (43).Therefore, in a second analysis, we also explored whether growth rates of *P. destructans* on individual female and male bats that were recaptured over winter clearly differed (43). For example, if one sex had higher fungal loads than the other and subsequently died of their infections, these highly infected bats will be omitted from the late hibernation population-level sample (77). By comparing the difference between late and early hibernation fungal loads on individual recaptured bats, we can directly assess whether female and male bats have different growth rates and better identify the timing of transmission differences that may underlie sex-biases in infection. Therefore, we compared the over winter increase in fungal loads on recaptured individuals (the log_10_ difference between late hibernation and early hibernation loads) using a LMM with sex as a fixed effect and site as a random effect (Model S1A, Appendix 5.1.1). Lastly, to explore differences in sex-based ecology that could lead to differences in infection between sexes, we also examined models including *a priori* hypothesized effects of roosting temperature and body mass on early hibernation prevalence and fungal loads but found no clear improvements over models with sex and species using AIC (Appendix 5.2 and 5.3). We used site as a random effect for all infection analyses and confirmed the performance of our infection models using 5-fold cross validation (see Supplemental Material for description).

#### ii. Individual survival

We examined differences between sexes in over winter survival using data from individual little brown bats, which were abundant enough to obtain reasonable sample sizes. We compared the probability of observing bats in late hibernation (March) that we observed in early hibernation (November). Bats affected by WNS often emerge prematurely (mid-winter) and die on the landscape (80), enabling the use of recapture as an estimate of apparent survival between sampling dates (79, 81). We used a mixed effects model with site as a random effect with a binomial distribution and a logit link to quantify how sex affected the probability of a bat being resighted over winter (Model 3A; Appendix 3.1).

#### iii. Population-level impact of disease

We estimated the proportion of female little brown bats sampled at each visit from a core set of sites (N=14) that were sampled in consecutive years to evaluate how female proportions were changing over time. We estimated the probability of sampling a female vs a male (1|0) over time using generalized linear mixed model with a binomial distribution and a logit link with continuous years since *P. destructans* invasion as a fixed effect, and site as a random effect (Model 3B; Appendix 3.2).

#### iv. RFID activity

To explore differences in autumn activity of female and male bats, we used a generalized linear mixed model with a binomial distribution and a logit link with sex and site as fixed effects (we did not include site as a random effect due to the small number of sites) and individual bat identification as a random effect (Model 4; Appendix 4.1). For our response variable, we treated each bat as a series of binomial trials where it could be detected (=1) or not detected (=0) on a given night throughout autumn swarm that any RFID system was operational at an entrance, as determined by the detection of a programmed test tag. We determined the period of autumn swarm to be between the dates in which more than 95% of all individual detections occurred (beginning of swarm; August 19) and all individual detections concluded (end of swarm; October 01). Results were qualitatively similar if we included all dates in which any bat was detected from early August to November. Since bats are nocturnal and daily activity patterns span two calendar days, we treated detections that occurred between hours 0:00-07:00 as a detection during the previous night for consistency with bat ecology. We used 5-fold cross-validation to check the performance of our model (see Supplemental Material for description).

## 3. RESULTS

We examined the effects of sex on *P. destructans* infections of five bat species during winter (Supplemental Table 1). We sampled a total of 665 females and 1071 males across all species, sites, years, and seasons (Supplemental Table 2A-D). On average, females had higher *P. destructans* prevalence than male bats in early winter (GLMM with site and species as random effects; female coef +/- SE: 0.630 +/- 0.221, P = 0.0043) and suffered from higher fungal loads (LMM with site and species as random effects; female coef +/- SE: 0.487 +/- 0.086, P < 0.0001). The model predicting early winter *P. destructans* prevalence, including species and an additive effect of sex, was better supported over the interactive version (ΔAIC = −3.11; Appendix 1.2-1.3), suggesting female infections are generally higher across all bat species in our study (GLMM with site as random effect; female coef +/- SE: 0.637 +/- 0.223, P = 0.0044; Figure 1; Appendix 1.1).

**Figure 1.**
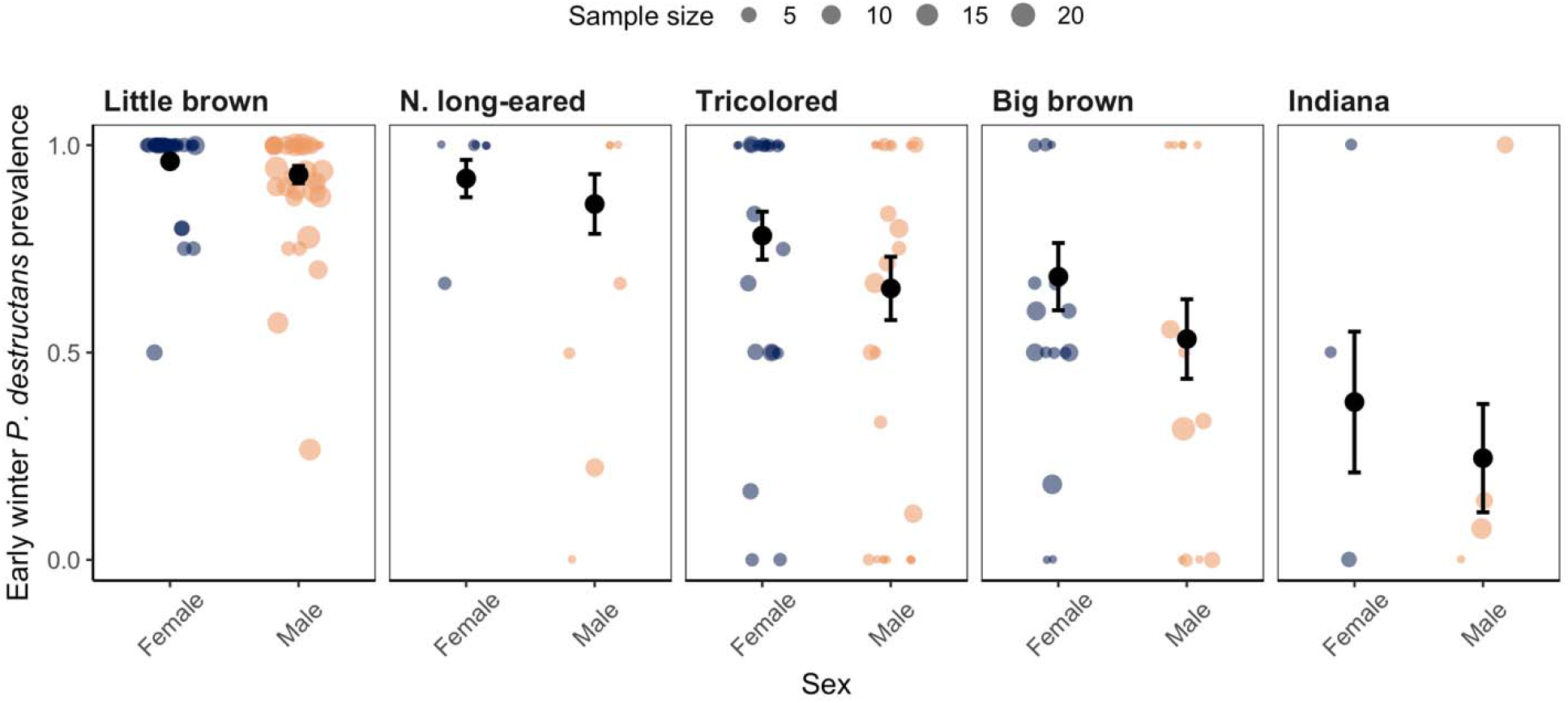
Early winter *P. destructans* prevalence by sex for five bat species. The colored points show the proportion of individuals that were infected at a given site and year. The size of colored points is scaled to the number of bats sampled during the site visit. The black circles show model predicted prevalence with standard error bars. Sample sizes are provided in Supplemental Table 2A.

The differences in infection prevalence during early winter aligned with higher fungal loads on female bats during the hibernation season. Across species, females began hibernation with higher fungal loads than males (LMM with site as random effect; female coef +/- SE: 0.490 +/- 0.093, P < 0.0001; Figure 2; Appendix 2.1). Fungal loads appeared to become more similar between females and males by the end of winter (smaller date slope for females (=0.405) vs males (=0.501); Figure 2; Appendix 2.4). However, this may be an artefact of higher female mortality (Figure 3; Appendix 3.1) such that individuals with higher fungal loads in early hibernation are less likely to survive to late winter, resulting in only bats with low early hibernation infections surviving to be sampled in late hibernation (77, 79). To assess whether mortality bias might be responsible for the apparent similarity of infections between sexes in late hibernation, we used data on differences in fungal loads of recaptured individual little brown bats sampled in both early and late winter. We found that the change in fungal loads over winter did not differ between sexes in the absence of selective mortality (LMM with site as a random effect; female coef +/- SE: 0.090 +/- 0.118, P = 0.448; Supplemental Figure1A; Appendix 5.1.1).

**Figure 2.**
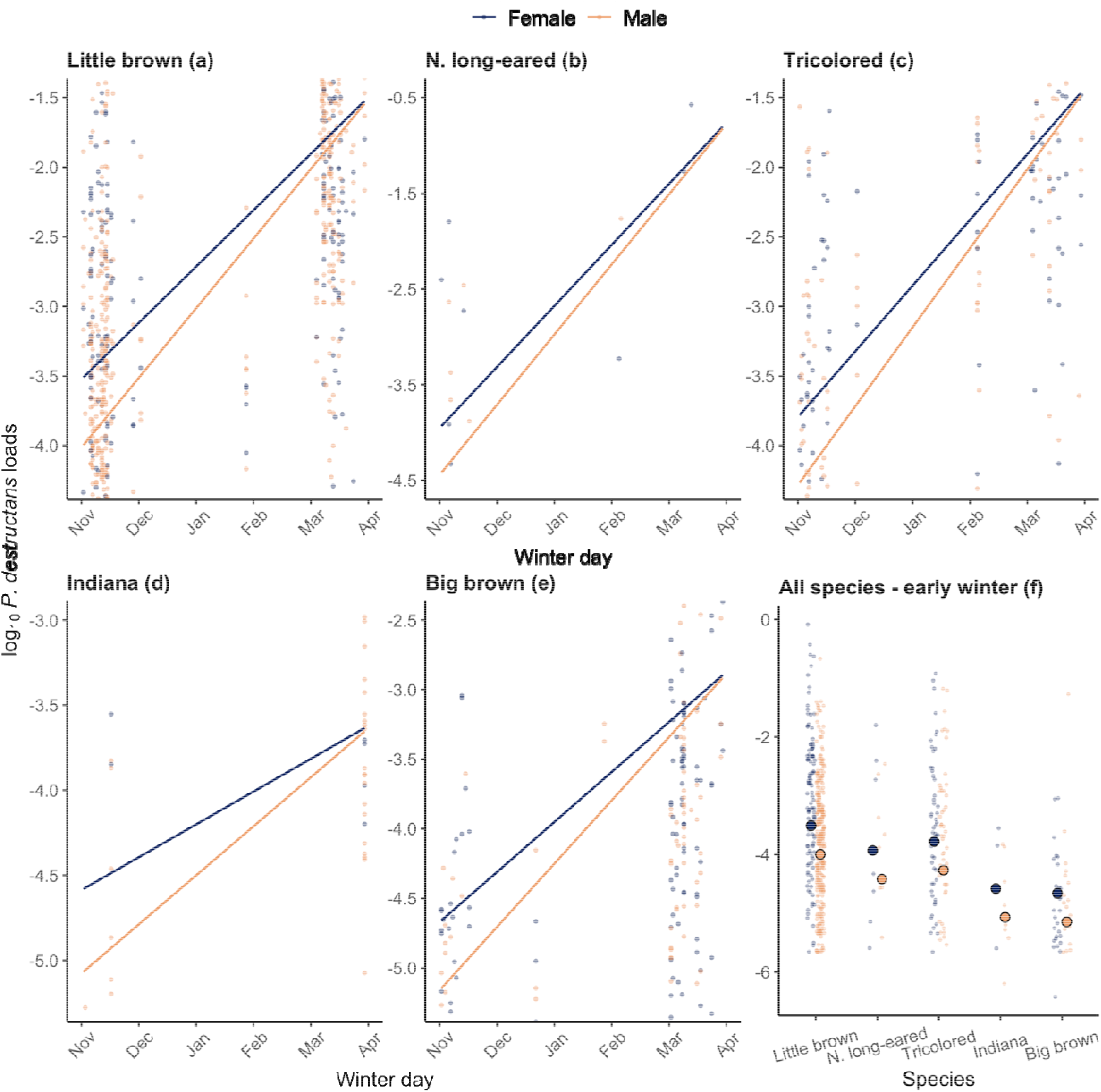
**(a-e)** Change in *P. destructans* infection severity (fungal loads) throughout winter by sex for each of the five host species. A point represents an individual bat, and females and males are shown in blue and orange, respectively. Lines represent model predicted fungal loads over time. Sample sizes by species are provided in Supplemental Table 2A. **(f)** Fungal loads in early winter by sex for each host species. Larger circles outlined in black show estimated fungal loads on November 1 extracted from model predictions used in a-e to visualize the differences in early winter infections across species.

**Figure 3.**
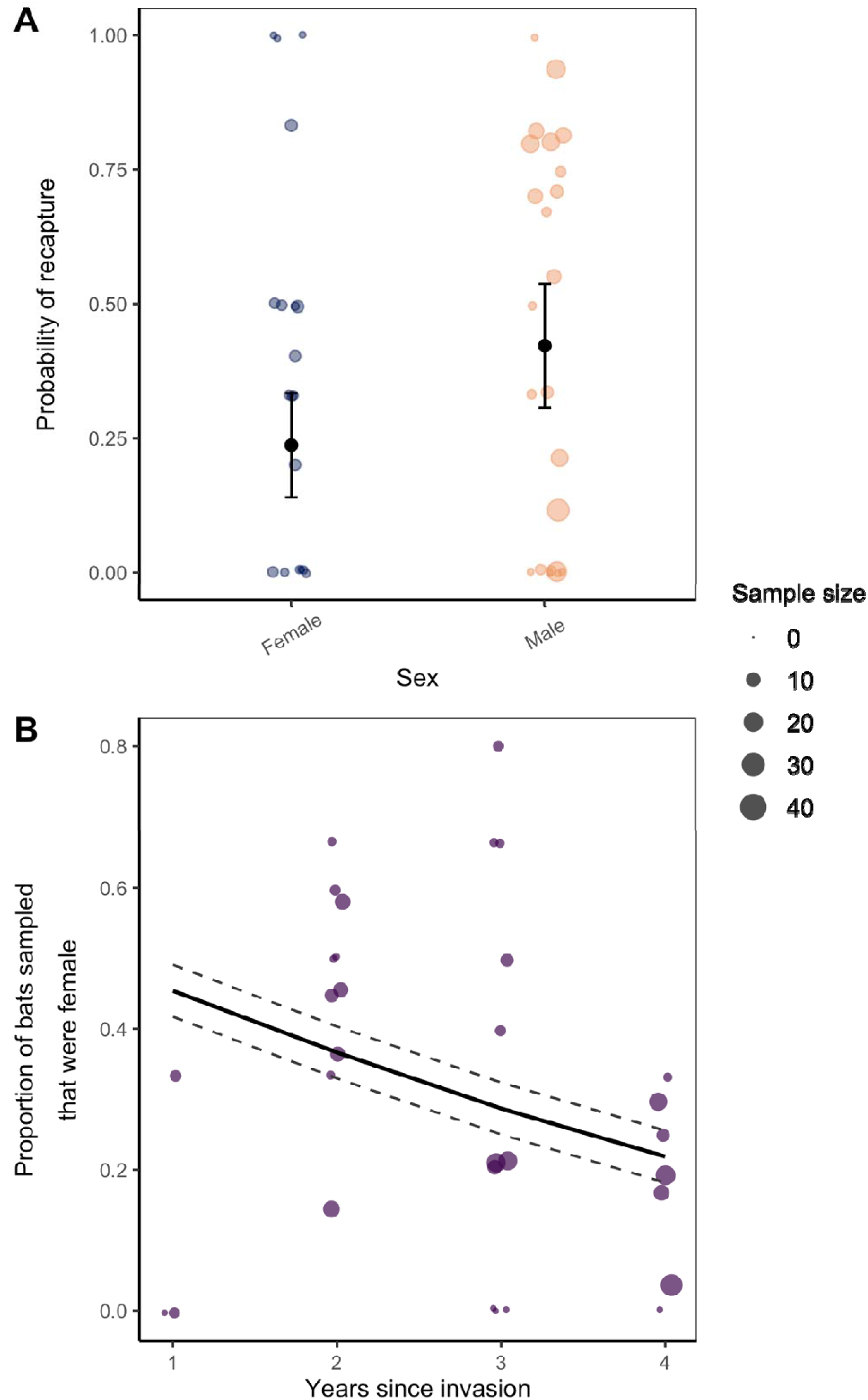
**(A)** Recapture probability of little brown bats by sex. Blue and orange points show the observed probability of recapture from early (November) and late (March) winter from a given site and year for females and males, respectively. Black points and vertical bars represent the estimated mean probability of recapture and standard errors. Sample sizes are provided in Supplemental Table 2B. **(B)** Proportional change in females with time since *P. destructans* invasion. Points show the proportion of all sampled individuals that were female at the same sites sampled over time since the invasion of *P. destructans*. The solid black line shows the model predicted proportion of females and dashed lines show the standard error around the model estimates. The size of points in both panels is scaled to the number of bats sampled during the site visit. Sample sizes are provided in Supplemental Table 2C.

We did not find support that hibernation temperatures or body mass significantly improved models of prevalence and fungal loads that already included sex (ΔAIC < 2; Appendix 5.2 and 5.3) and hibernation temperatures of male and female bats broadly overlapped (Supplemental Figure 2; Appendix 5.2).

At an individual scale, using data from little brown bats that were banded in early hibernation (Supplemental Table 2B), we found clear support that female little brown bats were less likely to be recaptured in late winter than males (GLMM with site as random effect; female coef +/- SE: - 0.855 +/- 0.380, P = 0.0240; Figure 3A; Appendix 3.1). At a population level, the proportion of females sampled in the same populations at the end of winter (Supplemental Table 2C) decreased with time since WNS invasion (GLMM with site as random effect; years since invasion coef +/- SE: −0.363 +/- 0.173, P = 0.0365; Figure 3B; Appendix 3.2). Collectively, these data suggest that females have higher, more severe infections (Figure 1, 2), and correspondingly experience higher mortality during winter (Figure 3).

Females begin hibernation with more severe infections than male bats and differential activity between sexes may influence infections. We found that the probability of a female being active on a given night was four-fold lower than males during autumn. (GLMM with individual bat as random effect; female coef +/- SE: −1.401 +/- 0.171, P < 0.0001; Figure 4; Supplemental Table 2D; Appendix 4.1).

**Figure 4.**
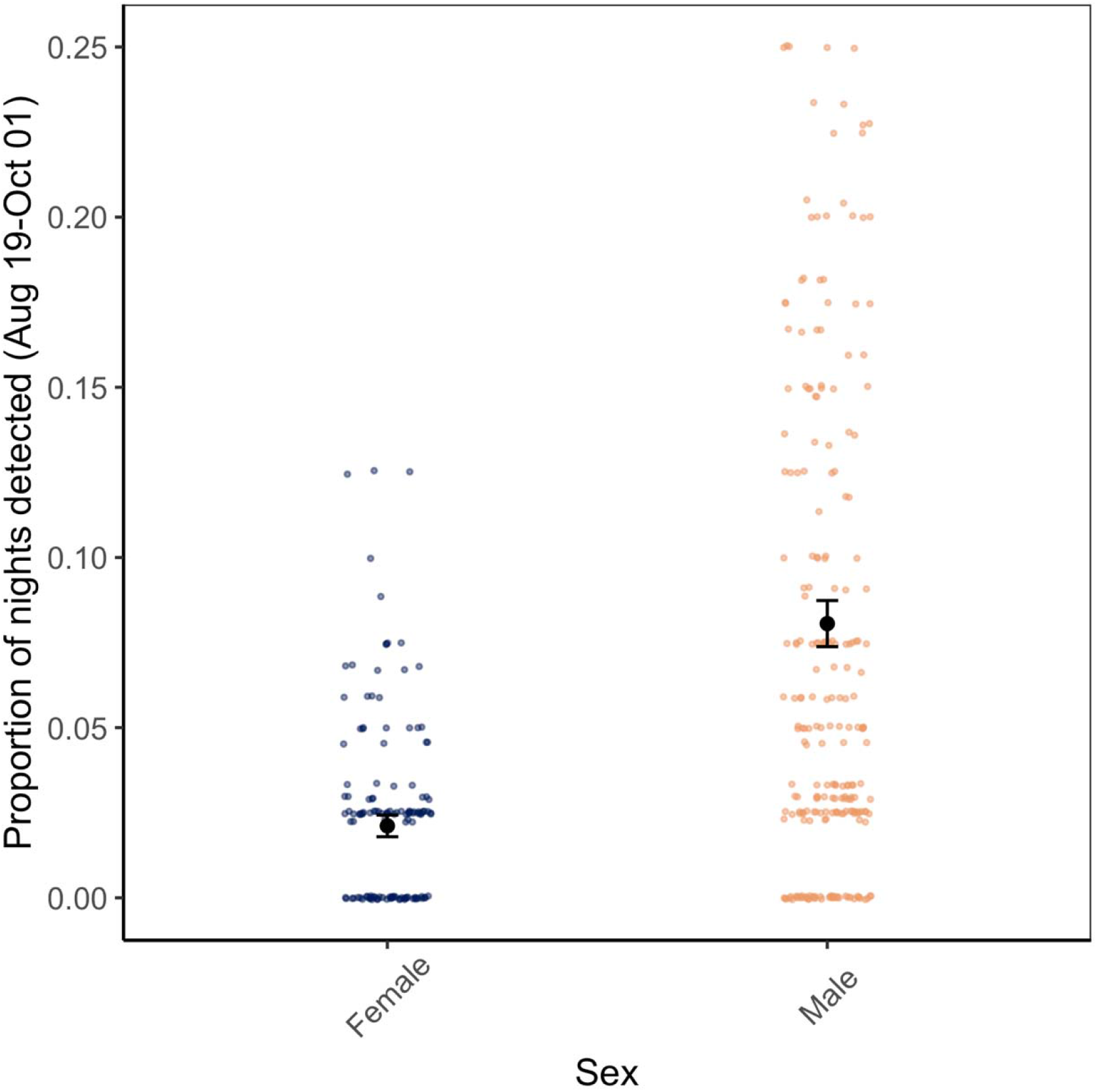
Differences in autumn activity of female and male little brown bats at three hibernacula. Each data point represents the proportion of nights an individual bat was active at a site. The proportion of nights active was calculated as the number of nights an individual was detected at least once on the RFID systems divided by the total number of nights the RFID systems at each site were operating throughout autumn swarm. Points at 0 represent tagged bats that were never detected on a reader. Black points show model predicted activity by sex with standard errors denoted with vertical bars. Sample sizes are provided in Supplemental Table 2D.

## 4. DISCUSSION

Our results demonstrate that sex differences in infection have an important role in disease dynamics. Across all species, females experienced higher infection prevalence (Figure 1) and fungal loads by the beginning of winter hibernation (Figure 2). In little brown bats, females were also less likely to be recaptured over winter (Figure 3A), as we would expect if more severe infections resulted in higher mortality among females (43, 56, 76, 77, 79). Finally, female-biased mortality associated with time since disease arrival likely contributed to population-level reductions in females over time (Figure 3B). Higher impacts to female bats may mask even more severe long-term impacts from WNS (46, 82) as a disproportionate loss of recruiting females may be expected to have more negative population-level consequences than male-biased declines.

We consistently observed higher pathogen prevalence and intensity in females than males across bat species (Figure 1, Figure 2). In other vertebrates, males maximize fitness at the cost of greater exposure and susceptibility to pathogens (33, 83–85), in part through higher testosterone concentrations which often correlate negatively with immune defense (86–88). Given male bats undergo their most substantial investment in reproduction during autumn swarm compared to females that invest relatively little towards reproduction throughout autumn and over winter (70), we expected infection to be higher in males. However, we found the opposite pattern with females having more severe disease. Drivers of higher female infections likely include sex-based physiological differences (25). Temperate female bats use fat stores acquired in autumn more slowly than males suggesting a more substantial constraint on energy expenditure compared to males throughout the autumn to spring period (66, 67). Therefore, different physiological strategies (e.g., torpor patterns) between female and male bats could be shaping sex-biased infections.

Differences in sex-based physiology and behavior were supported by our data showing that the proportion of nights males were active during autumn was substantially higher than females (Figure 4). Differences in autumn activity patterns are likely motivated by differences in reproductive energy allocation between sexes. For males, their fitness is enhanced by remaining active so they can maximize mating opportunities with females. However, female bats, which store sperm and delay ovulation until spring, may prefer to conserve energetic resources during fall and use torpor more extensively (67). As bats arrive to contaminated hibernacula (56, 58) for autumn swarm and are exposed to the pathogen, torpor use in autumn may permit pathogen growth as bats reduce their body temperatures to be within the thermal range for *P. destructans* replication. The relationship between sex-based activity and fungal intensity may arise through two potential pathways associated with torpor expression. First, less active females may provide favorable conditions for pathogen growth for a longer period prior to hibernation compared to males, thus resulting in the more severe female infections. Second, more active males may be able to actively inhibit pathogen growth through euthermia compared to females, as euthermic mammals mount more robust immune defenses than torpid or hibernating mammals (89, 90). Our activity estimates, which are consistent with other studies (65), suggest that vast differences between male and female bats’ energy use strategies during the mating period likely contribute to differences in infection.

We were less likely to recapture female little brown bats over winter than males, suggesting greater mortality in females (Figure 3A). Further, the sex ratio shifted to be more male-biased as WNS progressed (Figure 3B), suggesting that fungal infection may be driving female-specific mortality. Generally, infections increase on bats over winter before reaching a threshold at which bats die from their infections (43), and this bias in survival of the least infected individuals in early hibernation (Supplemental Figure 1B; Appendix 5.1.3) could explain why male and female infections become more similar at the end of hibernation (Figure 2). On average, female bats began hibernation with higher fungal loads than males (Figure 1) but have similar fungal growth rates (Supplemental Figure 1A; Appendix 5.1) suggesting that females reach high infection levels that result in mid-winter mortality, thus, are removed from the sample population by late winter. This relationship between high early hibernation fungal loads and mortality is further supported by previous studies that directly link early hibernation fungal loads with mortality and impacts (43, 79). Our findings are supported by previous work demonstrating that WNS severity is positively correlated with mortality, which is mechanistically linked by more frequent arousals (43, 47, 53, 77, 81). Several lines of evidence also suggest that females did not leave hibernation sites early and survive elsewhere. In our study region, emergence from hibernation in healthy populations does not begin until four to six weeks after our sampling (91). Given that the proportion of females declined with WNS progression (Figure 3B), our results suggest that disease associated mortality may be contributing to the reduction in overwinter recaptures of females. Previous studies on intersex differences in survival associated with WNS have found contrasting results. One study observed lower female survival in naturally infected bats (68) whereas another reported higher female survival in experimental infections where each sex was inoculated with the same dose of *P. destructans* at identical times (92). Our results suggest that differences in disease outcomes between females and males may originate from differences in pathogen growth during autumn. Therefore, inoculating both sexes with the same dose simultaneously in the experimental infection may have eliminated the underlying difference in early winter infections between female and males.

We provide the first spatiotemporally broad evidence from the field that sex-based survival of bats affected by WNS likely scales to population-level impacts. In the first year after *P. destructans* invasion, the mean proportion of females in late hibernation was 45%, but this percentage declined to 21% after four years of WNS (Figure 3B). The reduction in the percent of females with time since pathogen invasion differs from previous studies of healthy bat populations that show relatively consistent interannual proportions of females (93–96). This suggests that a comparable decline in females is not typical in non-diseased populations. Losing females at higher rates than males will likely affect how bat populations respond to WNS. First, reducing the number of females could limit the recruitment potential of bat populations and is likely to be especially critical for imperiled populations that remain small and vulnerable to Allee effects. Many temperate bat species, including little brown bats, form maternity colonies during summer when they cooperatively rear offspring and larger colony sizes afford females lower energy expenditure (97). If birth and offspring survival rates decline with density, as shown in other temperate bat species (98), the disproportionate loss of females could further impact recruitment. Second, distorted sex-ratios may negatively affect reproductive success during autumn. In other taxa, a significant increase in male to female sex-ratios resulted in increased stress to females from exacerbated pressure for males to mate with limited females, negatively affecting female fecundity (99, 100).

Our results include impacts to female bats up to four years after the arrival of WNS, which includes the epidemic phase of the disease, when the majority of mortality occurs (46). Females are likely under strong selection pressure to evolve mechanisms of survival given their increased mortality and will need to adapt for populations to rebound (101). Future work focusing on the effects of female infection and mortality biases on bat population persistence and recovery could benefit conservation efforts, especially as the negative effects are likely to compound over time if sex ratios become increasingly distorted.

We find a novel example in which female-biased infections may contribute to population-level impacts of an emerging disease. Our results provide a clear, yet less frequently observed, instance of an emerging pathogen that consistently impacts females more than males regardless of host species. We describe a new mechanistic explanation to female-biased infections that links temperature-dependent fungal growth to sex-specific seasonal physiology. Ultimately, disparate infections among demographic classes of hosts are fundamental for understanding and managing emerging infectious diseases, and cross-scale analyses can provide insights into the important consequences of demographic biases on disease systems.

## ACKNOWLEDGEMENTS

We thank Steffany Yamada for data curation support, Skylar Hopkins for analysis support, Rick Reynolds and Carl Herzog for logistical support, and the many landowners for site access.

## DISCLAIMER

Any use of trade, firm, or product names is for descriptive purposes only and does not imply endorsement by the U.S. Government.

## AUTHOR CONTRIBUTIONS

M.J.K.: conceptualization, investigation, methodology, visualization, data curation, writing-original draft, writing-review and editing, formal analysis; J.R.H.: conceptualization, investigation, methodology, visualization, funding acquisition, resources, project administration, data curation, writing-review and editing; J.P.W.: investigation, writing-review and editing; H. M.K.: investigation, writing-review and editing; J.A.R.: investigation; A.E.L.: resources, data curation, writing-review and editing; T.E.R.: resources, writing-review and editing; J.E.D.: investigation, writing-review and editing; W.H.S.: investigation; K.L.P.: investigation, writing-review and editing; J.T.F.: investigation, funding acquisition, writing-review and editing; A.M.K.: investigation, funding acquisition, writing-review and editing; K.E.L.: conceptualization, investigation, visualization, methodology, funding acquisition, resources, project administration, data curation, writing-original draft, writing-review and editing, supervision

All authors gave final approval for publication and agreed to be held accountable for the work performed therein.

## DATA, CODE, AND MATERIALS

The datasets and code generated in this study have been uploaded as electronic supporting material for review and will be deposited in Dryad Digital Repository upon final submission. Exact site locations are not disclosed to protect endangered species and landowners.

## ETHICS

All bat handling procedures were reviewed and approved by Virginia Tech Institute for Animal Care and Use Committee protocol 17-180.

