## Supplemental Table 1; Supplemental Figures for "Sex-biased infections scale to population impacts for an emerging wildlife disease"

### SUPPLEMENTAL MATERIALS

**Supplemental Table 1.** Summary of figures, statistical analyses, and the corresponding results throughout the manuscript.

| Figure; Model | Distribution | Response Variable | Fixed effect | Random effect | Data | Experimental unit | Findings |
|---|---|---|---|---|---|---|---|
| Fig. 1; Model 1 | Binomial | Pathogen detection on a bat (0\|1) in early hibernation | Sex+Species | Site | Infection (Supp Table 2A) | Individual swab | Across species, female bats have higher prevalence in early hibernation than males. |
| Fig. 2; Model 2 | Gaussian | Fungal loads (log10 ng DNA) on a bat | Sex*Date+Species*Date | Site | Infection (Supp Table 2A) | Individual swab | Across species, female bats begin hibernation at higher fungal loads, but loads become more similar by late hibernation at a population-level. |
| Fig. 3A; Model 3A | Binomial | Overwinter recapture (0\|1) | Sex | Site | Banded bats (Supp Table 2B) | Individual swab | Female little brown bats are less likely to survive over winter than males. |
| Fig. 3B; Model 3B | Binomial | Sex of bat sampled (0 = male\|1 = female) | Years since pathogen invasion (YSW; continuous) | Site | Infection (Supp Table 2C) | Individual bat | The probability of sampling a female decreased with years since *P. destructans* invasion. |
| Fig. 4; Model 4 | Binomial | Bat detected (active = 1) or not (inactive = 0) | Sex + Site | Individual | Fall activity (Supp Table 2D) | Individual detection | Female bats spend fewer nights active than males throughout autumn. |

| | | | | | | | |
|---|---|---|---|---|---|---|---|
| Supp Fig. 1A; Model S1A | Gaussian | Overwinter increase in fungal load | Sex | Site | Banded bats that survived over winter (Supp Table 2B) | A change in fungal load value from an individual bat captured in both early and late hibernation. | Fungal growth rates overwinter are similar between males and females that survive WNS. |

**Supplemental Figures**

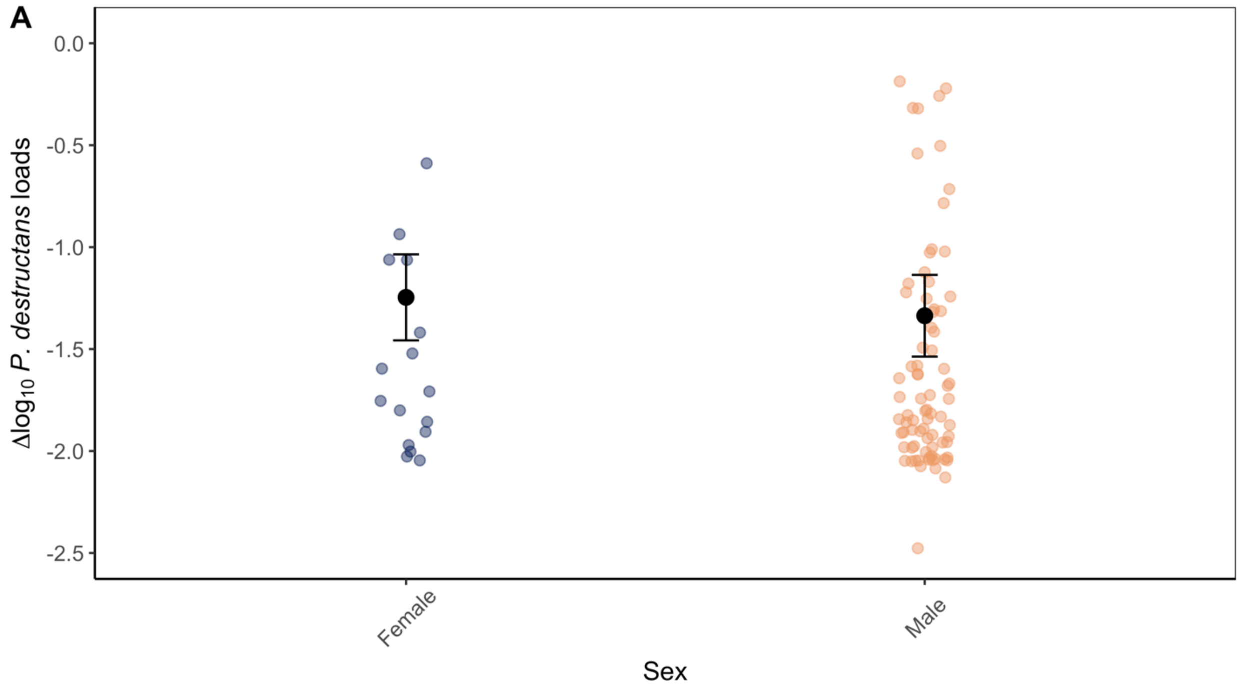


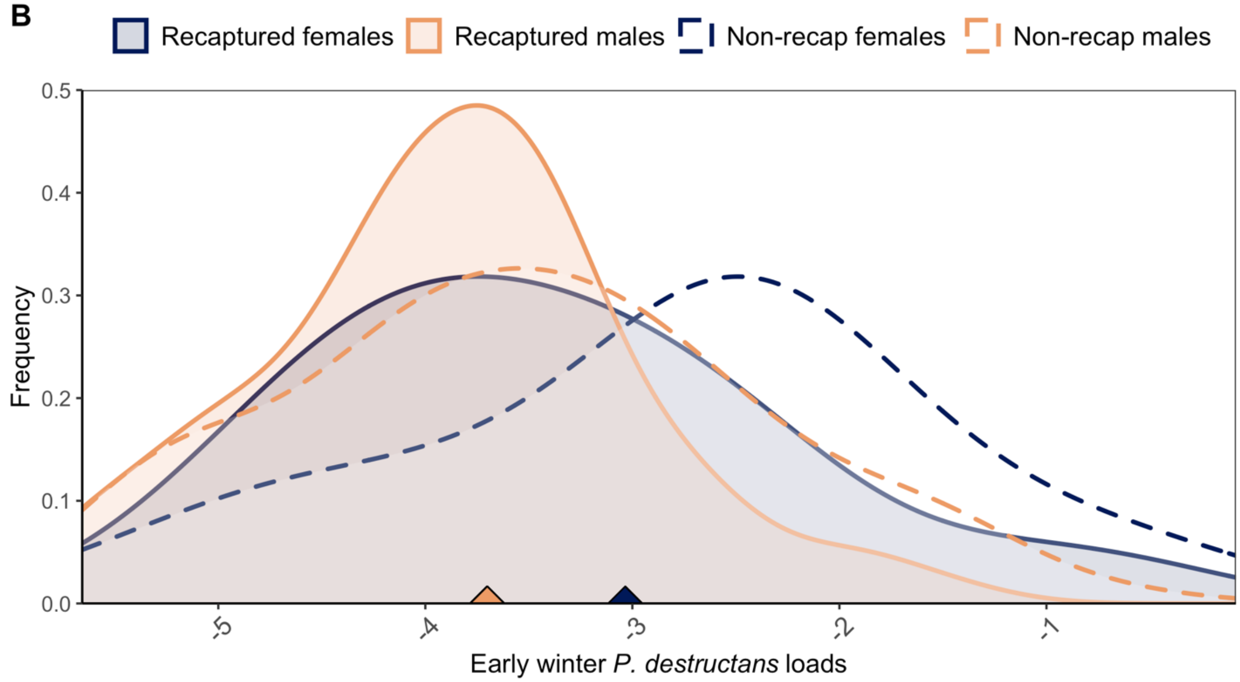


**Supplemental Figure 1: (A)** Change in over winter fungal loads of recaptured little brown bats by sex. Increase in pathogen loads over winter on female and male bats that survived their infections were similar, suggesting that higher early winter infections contribute to lower female survival as opposed to differential pathogen growth rates between sexes over winter. Black points are mean estimates from Model S1A with standard error bars. **(B)** Density plot of the relationship between early winter fungal loads and recapture category of females in blue and males in orange. Solid lines show the frequency of observed recaptured individuals and dashed lines show the frequencies of individuals that were not recaptured. Along the x-axis, solid triangles denote the observed mean early winter fungal loads of females and males.

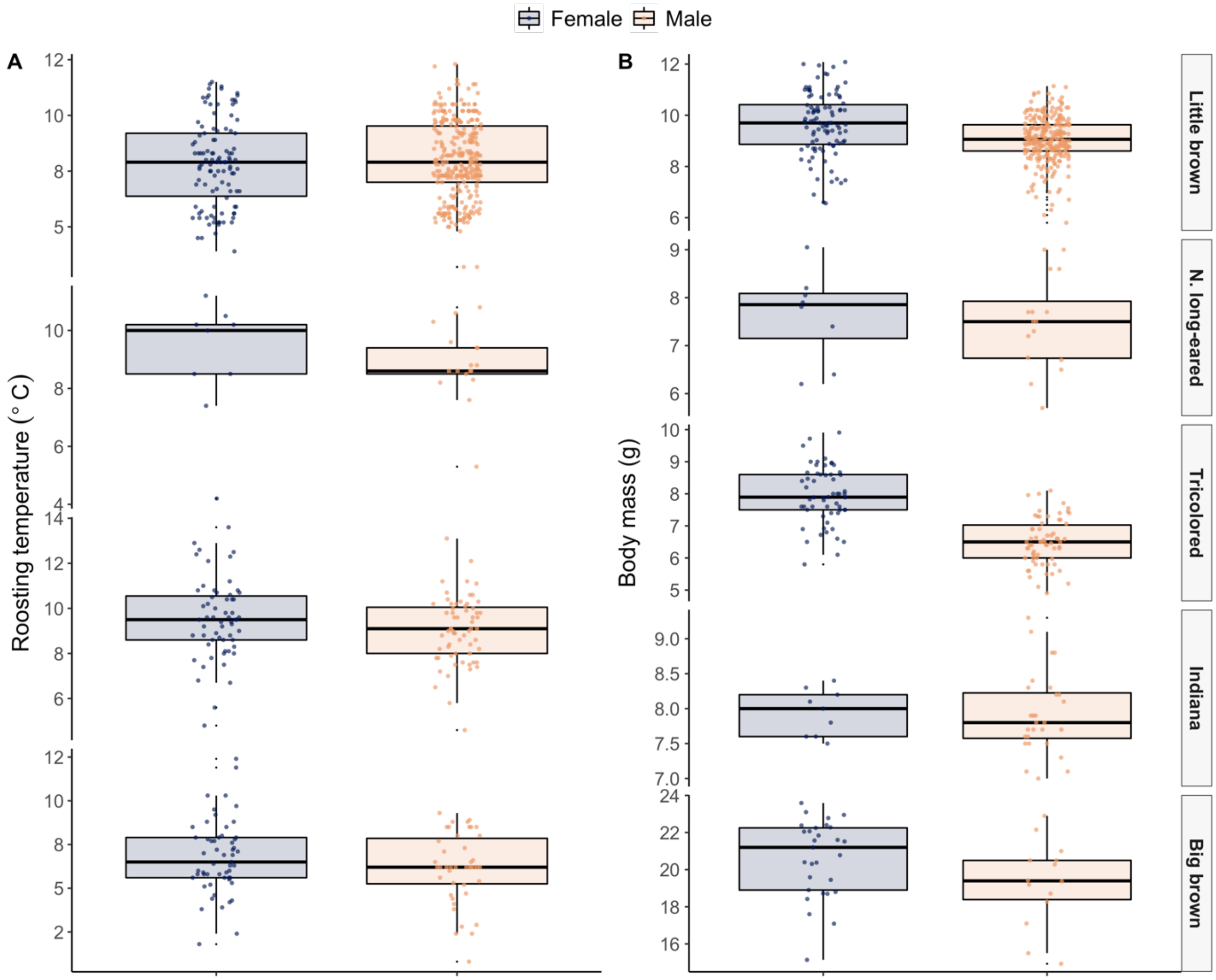


**Supplemental Figure 2.** Early hibernation roosting temperature (**A; left**) and body mass (**B; right**) of four species for which there were sufficient data to analyze the two traits between sexes and on early hibernation infections. Indiana bats were not included due to data deficiency. Roosting temperature varied minimally between females and males of most species. Body masses were generally higher for females while temperatures were generally lower, but neither variable improved model predictions of disease over the base models (Model 1 or Model 2; Appendix 5.2 and 5.3).

**Model performance checks**

We included checks of our model performance using k-fold cross validation. For the logistic Model 1 (pd(0|1)~sex+species), we performed five-fold cross validation of 1000 random divisions of our dataset and calculated the average area under the curve for the resulting 1000 receiver operating characteristics (ROC). These results indicate that ~75% of our test data was successfully predicted by the training models. We performed five-fold cross validation of Model 2 (fungal.loads~sex*date+species*date), iteratively simulated 1000 random divisions of the dataset and calculated subsequent mean R2 and mean RMSE (mean R2 = 0.547; mean RMSE = 0.989). For models of individual bat recapture (Model 3A) and the proportion of females sampled in late hibernation (Model 3B), sample sizes were not sufficiently large enough to perform cross-validation, however all the 95% confidence intervals of the estimated marginal means (e.g. the coefs in 'response' space, model estimates) always included the observed means. We also performed 5-fold cross validation on the autumn activity analysis, Model 4 (active(0|1)~sex+site), using the same approach as we did with the logistic Model 1 above. These results indicated that ~74% of our test data was successfully predicted by the training models.
