## Appendix; Supplemental Table 2A-2D for "Sex-biased infections scale to population impacts for an emerging wildlife disease": kailingetal_sexbias_pdinf_appendix_rev1.html

Appendix for Kailing et al. Sex biased Pd Infection Ms


### Appendix for Kailing et al. Sex biased Pd Infection Ms

- 1 How does **prevalence
  (gd)** differ by sex in early hibernation?
  - 1.1 Model 1: Species specific
    differences by sex - sex and species as fixed effects; site as random
    effect
  - 1.2 Check performance of Model 1
    against other model structures. All models include site as a random
    effect.
  - 1.3 Check support for interaction
    between sex and species
  - 1.4
    ***Result:***
  - 1.5 View prevalence plot (Figure
    1)
- 2 How does **infection severity
  (fungal loads; lgdL)** differ by host sex?
  - 2.1 Model 2: Population-level sex
    differences in fungal loads by species *over winter* - sex,
    species, and date as fixed effects; site as random effect
  - 2.2 Check performance of Model 2
    against other model structures. All models include site as a random
    effect.
  - 2.3
    ***Result:***
  - 2.4 View fungal loads plot (Figure
    2a-f)
- 3 Do sex-biased infections affect
  **survival** or **population structure**?
  - 3.1 Model 3A: Effect of host sex on
    probability of re-sighting (e.g., apparent survival) - sex as fixed
    effect; site as random effect
    - 3.1.1 Check performance of Model 3A
      compared to null. Both models include site as a random effect.
    - 3.1.2
      ***Result:***
  - 3.2 Model 3B: Effect of years since
    pathogen invasion on the proportion of bats that are female in late
    hibernation - years since disease invasion as fixed effect; site as
    random effect
    - 3.2.1 Check performance of Model3B
      compared to null. Both models include site as a random effect.
    - 3.2.2
      ***Result:***
  - 3.3 View little brown bat impacts plot
    (Figure 3A-B)
- 4 Does **autumn swarm
  activity** differ by sex?
  - 4.1 Model 4: Sex differences in autumn
    swarm - sex and site as fixed effects; individual ID as random
    effect
  - 4.2 Check performance of Model 4
    compared to other model structures.
  - 4.3
    ***Result:***
  - 4.4 View autumn activity plot (Figure
    4)
- 5 Supplemental analyses and
  figures
  - 5.1 Explore relationships between
    survival and fungal loads by sex
    - 5.1.1 Model S1A: Do female and male
      **changes in fungal loads** differ over winter? sex as
      fixed effect; site as random effect
    - 5.1.2 View change in fungal loads plot
      (Supp Figure 1A)
    - 5.1.3 Model S1B: What is the
      relationship between **early winter loads and recapture
      frequency** of females and males? sex and recapture status as
      fixed effects
    - 5.1.4 View plot of recapture status
      and sex over early hibernation fungal loads (Supp Figure 1B)
    - 5.1.5
      ***Result:***
  - 5.2 Do differences in early
    hibernation roosting temperature explain female biased infections? (Supp
    Fig 2A)
    - 5.2.1 Are there sex differences in
      roosting temperature among species?
      - 5.2.1.1 View model predicted means
        (emmean) and standard errors (SE) by sex and species
      - 5.2.1.2
        ***Result:***
    - 5.2.2 Explore whether addition of
      temperature term in prevalence model improves predictions.
      - 5.2.2.1 Early hibernation
        **prevalence** model comparison to models with temperature.
        All models have site as random effect.
      - 5.2.2.2
        ***Result:***
    - 5.2.3 Now, explore whether addition of
      temperature term in fungal loads model improves estimates.
      - 5.2.3.1 Early hibernation
        **fungal load** model comparison with temperature models.
        All models have site as random effect.
      - 5.2.3.2 Check whether there is support
        for temperature term (fl2 = sex+species+temp)
      - 5.2.3.3
        ***Result:***
  - 5.3 Do differences in early
    hibernation **body mass** explain female biased infections?
    (Supp Fig 2B)
    - 5.3.1 Early hibernation
      **prevalence** model comparisons to models with body mass
      terms. All models have site as a random effect.
    - 5.3.2 Early hibernation **fungal
      loads** model comparisons to models with body mass terms. All
      models have site as a random effect.
    - 5.3.3
      ***Result:***
  - 5.4 View plots of early hibernation
    roosting temperature and mass across species (Supp Fig 2)

**OVERVIEW:**

Each model is parameterized in relation to the reference level or
intercept (F = Female; MYLU = little brown bat) and each parameter
estimate represents the difference from this reference level,
incorporating the effect of each fixed effect (i.e., species, sex, date,
etc). An example in Model 1: the “Estimate” for sexM listed in the
output summary is the inverse logit fraction of the male little brown
bat population infected relative to females.

### 1 How does **prevalence (gd)** differ by sex in early hibernation?

#### 1.1 Model 1: Species specific differences by sex - sex and species as fixed effects; site as random effect

```
Generalized linear mixed model fit by maximum likelihood (Laplace
  Approximation) [glmerMod]
 Family: binomial  ( logit )
Formula: gd ~ species + sex + (1 | site)
   Data: est.dat
Control: glmerControl(optimizer = "bobyqa", optCtrl = list(maxfun = 1e+05))
 Subset: season2 == "h.earl"

     AIC      BIC   logLik deviance df.resid 
   694.2    727.2   -340.1    680.2      813 

Scaled residuals: 
     Min       1Q   Median       3Q      Max 
-10.7876  -0.2530   0.2482   0.4296   2.8296 

Random effects:
 Groups Name        Variance Std.Dev.
 site   (Intercept) 1.463    1.209   
Number of obs: 820, groups:  site, 34

Fixed effects:
            Estimate Std. Error z value Pr(>|z|)    
(Intercept)   3.2105     0.3655   8.784  < 2e-16 ***
speciesMYSE  -0.7737     0.5323  -1.454  0.14606    
speciesPESU  -1.9343     0.3013  -6.419 1.37e-10 ***
speciesMYSO  -3.6971     0.7366  -5.019 5.19e-07 ***
speciesEPFU  -2.4431     0.3239  -7.544 4.56e-14 ***
sexM         -0.6370     0.2234  -2.851  0.00435 ** 
---
Signif. codes:  0 '***' 0.001 '**' 0.01 '*' 0.05 '.' 0.1 ' ' 1

Correlation of Fixed Effects:
            (Intr) spMYSE spPESU spMYSO spEPFU
speciesMYSE -0.116                            
speciesPESU -0.496  0.175                     
speciesMYSO -0.292  0.038  0.195              
speciesEPFU -0.415  0.098  0.346  0.112       
sexM        -0.510  0.047  0.240  0.036  0.284
```

#### 1.2 Check performance of Model 1 against other model structures. All models include site as a random effect.

- ‘null’ model: gd~1

```
|   |Modnames                 |  K|      AIC| Delta_AIC|  ModelLik|     AICWt|        LL|    Cum.Wt|
|:--|:------------------------|--:|--------:|---------:|---------:|---------:|---------:|---------:|
|4  |species+sex              |  7| 694.2394|  0.000000| 1.0000000| 0.7948148| -340.1197| 0.7948148|
|5  |species * sex            | 11| 697.3470|  3.107562| 0.2114470| 0.1680612| -337.6735| 0.9628760|
|3  |species                  |  6| 700.5184|  6.278981| 0.0433048| 0.0344193| -344.2592| 0.9972953|
|2  |sex + random eff:species |  4| 705.6056| 11.366238| 0.0034029| 0.0027047| -348.8028| 1.0000000|
|1  |null                     |  2| 785.5326| 91.293252| 0.0000000| 0.0000000| -390.7663| 1.0000000|
```

#### 1.3 Check support for interaction between sex and species

- prA: sex+species
- prB: sex\*species

```
Data: est.dat
Subset: season2 == "h.earl"
Models:
pr.null: gd ~ 1 + (1 | site)
prA: gd ~ species + sex + (1 | site)
prB: gd ~ species * sex + (1 | site)
        npar    AIC    BIC  logLik deviance    Chisq Df Pr(>Chisq)    
pr.null    2 785.53 794.95 -390.77   781.53                           
prA        7 694.24 727.20 -340.12   680.24 101.2933  5     <2e-16 ***
prB       11 697.35 749.15 -337.67   675.35   4.8924  4     0.2985    
---
Signif. codes:  0 '***' 0.001 '**' 0.01 '*' 0.05 '.' 0.1 ' ' 1
```

#### 1.4 ***Result:***

- ∆AIC of species + sex model compared to other models is
  **>-3.1**. Reported model is best supported.
- There is no clear support for interaction between species and
  sex.
- Across species, female bats have higher prevalence in early
  hibernation than males.

#### 1.5 View prevalence plot (Figure 1)

### 2 How does **infection severity (fungal loads; lgdL)** differ by host sex?

#### 2.1 Model 2: Population-level sex differences in fungal loads by species *over winter* - sex, species, and date as fixed effects; site as random effect

- ‘*pdate.scale*’ is a date variable converted to monthly units
  then scaled to start at Day 0.

```
Linear mixed model fit by REML ['lmerMod']
Formula: lgdL ~ species * pdate.scale + sex * pdate.scale + (1 | site)
   Data: est.dat

REML criterion at convergence: 4161.4

Scaled residuals: 
    Min      1Q  Median      3Q     Max 
-5.2371 -0.5644  0.0152  0.6149  3.8040 

Random effects:
 Groups   Name        Variance Std.Dev.
 site     (Intercept) 0.3644   0.6037  
 Residual             0.9223   0.9603  
Number of obs: 1463, groups:  site, 42

Fixed effects:
                        Estimate Std. Error t value
(Intercept)             -3.52333    0.13261 -26.569
speciesMYSE             -0.42879    0.26808  -1.600
speciesPESU             -0.27203    0.12427  -2.189
speciesMYSO             -1.06358    0.33545  -3.171
speciesEPFU             -1.14643    0.18118  -6.328
pdate.scale              0.40523    0.02659  15.238
sexM                    -0.48994    0.09288  -5.275
speciesMYSE:pdate.scale  0.23282    0.13602   1.712
speciesPESU:pdate.scale  0.06707    0.03719   1.803
speciesMYSO:pdate.scale -0.21225    0.07366  -2.881
speciesEPFU:pdate.scale -0.04564    0.04777  -0.955
pdate.scale:sexM         0.09528    0.02704   3.523

Correlation of Fixed Effects:
            (Intr) spMYSE spPESU spMYSO spEPFU pdt.sc sexM   sMYSE: sPESU:
speciesMYSE -0.109                                                        
speciesPESU -0.301  0.096                                                 
speciesMYSO -0.078  0.033  0.076                                          
speciesEPFU -0.267  0.063  0.170  0.034                                   
pdate.scale -0.522  0.126  0.294  0.089  0.271                            
sexM        -0.495  0.065  0.172 -0.013  0.214  0.597                     
spcsMYSE:p.  0.066 -0.564 -0.036 -0.023 -0.044 -0.138 -0.035              
spcsPESU:p.  0.200 -0.068 -0.701 -0.047 -0.146 -0.471 -0.160  0.075       
spcsMYSO:p.  0.074 -0.035 -0.071 -0.862 -0.050 -0.162  0.000  0.037  0.112
spcsEPFU:p.  0.195 -0.056 -0.139 -0.029 -0.859 -0.364 -0.192  0.058  0.215
pdt.scl:sxM  0.404 -0.057 -0.160  0.009 -0.176 -0.739 -0.801  0.055  0.233
            sMYSO: sEPFU:
speciesMYSE              
speciesPESU              
speciesMYSO              
speciesEPFU              
pdate.scale              
sexM                     
spcsMYSE:p.              
spcsPESU:p.              
spcsMYSO:p.              
spcsEPFU:p.  0.067       
pdt.scl:sxM -0.007  0.230
```

- Calculate p-value from t-value for sex difference estimates

```
[1] 1.534453e-07
```

#### 2.2 Check performance of Model 2 against other model structures. All models include site as a random effect.

- ‘null’ model: lgdL~1

```
|   |Modnames                  |  K|      AIC| Delta_AIC| ModelLik|     AICWt|        LL|    Cum.Wt|
|:--|:-------------------------|--:|--------:|---------:|--------:|---------:|---------:|---------:|
|6  |species * date+sex * date | 14| 4146.966|    0.0000| 1.00e+00| 0.9999021| -2059.483| 0.9999021|
|5  |species+date+sex          |  9| 4165.429|   18.4635| 9.79e-05| 0.0000979| -2073.715| 1.0000000|
|4  |species+sex               |  8| 4903.466|  756.5003| 0.00e+00| 0.0000000| -2443.733| 1.0000000|
|2  |sex + random eff:species  |  5| 4916.087|  769.1211| 0.00e+00| 0.0000000| -2453.043| 1.0000000|
|3  |species                   |  7| 4918.534|  771.5681| 0.00e+00| 0.0000000| -2452.267| 1.0000000|
|1  |null                      |  3| 5037.020|  890.0540| 0.00e+00| 0.0000000| -2515.510| 1.0000000|
```

#### 2.3 ***Result:***

- ∆AIC of species \* date + sex \* date model compared to other models
  is **>-18.4**. Reported model is best supported.
- By early hibernation, females have more severe infections compared
  to males but appear to have lower fungal growth. However, this may be
  driven by a survival bias (see Supp Analysis 1; Figure S1A).

#### 2.4 View fungal loads plot (Figure 2a-f)

### 3 Do sex-biased infections affect **survival** or **population structure**?

#### 3.1 Model 3A: Effect of host sex on probability of re-sighting (e.g., apparent survival) - sex as fixed effect; site as random effect

```
Generalized linear mixed model fit by maximum likelihood (Laplace
  Approximation) [glmerMod]
 Family: binomial  ( logit )
Formula: RecapturedSameYearYN.x ~ sex.x + (1 | site2.x)
   Data: wr
Control: glmerControl(optimizer = "bobyqa", optCtrl = list(maxfun = 1e+05))

     AIC      BIC   logLik deviance df.resid 
   273.7    284.1   -133.8    267.7      237 

Scaled residuals: 
    Min      1Q  Median      3Q     Max 
-2.0670 -0.6478 -0.2928  0.7152  5.1007 

Random effects:
 Groups  Name        Variance Std.Dev.
 site2.x (Intercept) 1.915    1.384   
Number of obs: 240, groups:  site2.x, 13

Fixed effects:
            Estimate Std. Error z value Pr(>|z|)  
(Intercept)  -1.1691     0.5366  -2.179   0.0293 *
sex.xM        0.8548     0.3788   2.257   0.0240 *
---
Signif. codes:  0 '***' 0.001 '**' 0.01 '*' 0.05 '.' 0.1 ' ' 1

Correlation of Fixed Effects:
       (Intr)
sex.xM -0.514
```

##### 3.1.1 Check performance of Model 3A compared to null. Both models include site as a random effect.

- ‘null’ model: winter recap~1

```
|   |Modnames |  K|      AIC| Delta_AIC|  ModelLik|     AICWt|        LL|    Cum.Wt|
|:--|:--------|--:|--------:|---------:|---------:|---------:|---------:|---------:|
|2  |sex      |  3| 273.6612|  0.000000| 1.0000000| 0.8287891| -133.8306| 0.8287891|
|1  |null     |  2| 276.8154|  3.154139| 0.2065796| 0.1712109| -136.4077| 1.0000000|
```

##### 3.1.2 ***Result:***

- Including sex improves the ability to predict whether a bat will be
  re-sighted in late hibernation.
- Female little brown bats are less likely to survive over winter than
  males.

#### 3.2 Model 3B: Effect of years since pathogen invasion on the proportion of bats that are female in late hibernation - years since disease invasion as fixed effect; site as random effect

```
Generalized linear mixed model fit by maximum likelihood (Laplace
  Approximation) [glmerMod]
 Family: binomial  ( logit )
Formula: sex2 ~ ysw + (1 | site)
   Data: late
Control: glmerControl(optimizer = "bobyqa", optCtrl = list(maxfun = 1e+05))

     AIC      BIC   logLik deviance df.resid 
   295.0    305.5   -144.5    289.0      244 

Scaled residuals: 
    Min      1Q  Median      3Q     Max 
-0.9210 -0.6519 -0.5169  1.2315  2.2526 

Random effects:
 Groups Name        Variance Std.Dev.
 site   (Intercept) 0.101    0.3178  
Number of obs: 247, groups:  site, 14

Fixed effects:
            Estimate Std. Error z value Pr(>|z|)  
(Intercept)   0.1780     0.5078   0.350   0.7260  
ysw          -0.3628     0.1734  -2.092   0.0365 *
---
Signif. codes:  0 '***' 0.001 '**' 0.01 '*' 0.05 '.' 0.1 ' ' 1

Correlation of Fixed Effects:
    (Intr)
ysw -0.939
```

##### 3.2.1 Check performance of Model3B compared to null. Both models include site as a random effect.

- ‘null’ model: sex~1

```
|   |Modnames |  K|      AIC| Delta_AIC|  ModelLik|     AICWt|        LL|    Cum.Wt|
|:--|:--------|--:|--------:|---------:|---------:|---------:|---------:|---------:|
|2  |ysw      |  3| 295.0053|  0.000000| 1.0000000| 0.7351623| -144.5026| 0.7351623|
|1  |null     |  2| 297.0472|  2.041948| 0.3602439| 0.2648377| -146.5236| 1.0000000|
```

##### 3.2.2 ***Result:***

- Including years since white-nose syndrome invasion improves the
  ability to predict whether a bat sampled in late hibernation will be
  female.
- The probability of sampling a female decreased with years since
  *P. destructans* invasion.

#### 3.3 View little brown bat impacts plot (Figure 3A-B)

### 4 Does **autumn swarm activity** differ by sex?

#### 4.1 Model 4: Sex differences in autumn swarm - sex and site as fixed effects; individual ID as random effect

```
Generalized linear mixed model fit by maximum likelihood (Laplace
  Approximation) [glmerMod]
 Family: binomial  ( logit )
Formula: prop.act ~ sex + site2 + (1 | pit_id)
   Data: fall
Weights: rdr.nt
Control: glmerControl(optimizer = "bobyqa", optCtrl = list(maxfun = 1e+05))

     AIC      BIC   logLik deviance df.resid 
  1973.4   1993.6   -981.7   1963.4      420 

Scaled residuals: 
    Min      1Q  Median      3Q     Max 
-3.4381 -0.6279 -0.0167  0.3083  3.1567 

Random effects:
 Groups Name        Variance Std.Dev.
 pit_id (Intercept) 1.298    1.139   
Number of obs: 425, groups:  pit_id, 373

Fixed effects:
            Estimate Std. Error z value Pr(>|z|)    
(Intercept)  -4.1129     0.1618 -25.412  < 2e-16 ***
sexM          1.4007     0.1710   8.189 2.63e-16 ***
site2A MIE    1.1327     0.1795   6.311 2.77e-10 ***
site2DEN OC  -0.2998     0.1816  -1.651   0.0988 .  
---
Signif. codes:  0 '***' 0.001 '**' 0.01 '*' 0.05 '.' 0.1 ' ' 1

Correlation of Fixed Effects:
            (Intr) sexM   s2AMIE
sexM        -0.759              
site2A MIE  -0.243 -0.157       
site2DEN OC -0.362  0.018  0.319
```

#### 4.2 Check performance of Model 4 compared to other model structures.

- ‘null’ model: proportion of nights active~1.

```
|   |Modnames |  K|      AIC| Delta_AIC| ModelLik| AICWt|         LL| Cum.Wt|
|:--|:--------|--:|--------:|---------:|--------:|-----:|----------:|------:|
|3  |sex+site |  5| 1973.377|   0.00000|        1|     1|  -981.6887|      1|
|2  |sex      |  3| 2018.967|  45.58928|        0|     0| -1006.4834|      1|
|1  |null     |  2| 2093.959| 120.58183|        0|     0| -1044.9797|      1|
```

#### 4.3 ***Result:***

- The additive effects of sex and site is better supported over models
  including sex alone or the null model.
- Female bats spend fewer nights active than males throughout
  autumn.

#### 4.4 View autumn activity plot (Figure 4)

### 5 Supplemental analyses and figures

#### 5.1 Explore relationships between survival and fungal loads by sex

##### 5.1.1 Model S1A: Do female and male **changes in fungal loads** differ over winter? sex as fixed effect; site as random effect

```
Linear mixed model fit by REML ['lmerMod']
Formula: log_Dloads2 ~ sex.x + (1 | site2.x)
   Data: wr1

REML criterion at convergence: 130.3

Scaled residuals: 
    Min      1Q  Median      3Q     Max 
-3.4156 -0.5598 -0.1691  0.5704  2.9022 

Random effects:
 Groups   Name        Variance Std.Dev.
 site2.x  (Intercept) 0.3437   0.5862  
 Residual             0.1672   0.4090  
Number of obs: 97, groups:  site2.x, 10

Fixed effects:
            Estimate Std. Error t value
(Intercept) -1.24637    0.21081  -5.912
sex.xM      -0.09011    0.11819  -0.762

Correlation of Fixed Effects:
       (Intr)
sex.xM -0.372
```

- Calculate p-value from t-value for change in fungal loads
  estimates

```
[1] 0.4481692
```

##### 5.1.2 View change in fungal loads plot (Supp Figure 1A)

##### 5.1.3 Model S1B: What is the relationship between **early winter loads and recapture frequency** of females and males? sex and recapture status as fixed effects

```
Call:
lm(formula = lgdL.x ~ sex.x + RecapturedSameYearYN.x, data = wr)

Residuals:
     Min       1Q   Median       3Q      Max 
-2.78008 -0.68981  0.06046  0.65047  2.82307 

Coefficients:
                        Estimate Std. Error t value Pr(>|t|)    
(Intercept)              -2.8736     0.1511 -19.017  < 2e-16 ***
sex.xM                   -0.6194     0.1626  -3.810 0.000177 ***
RecapturedSameYearYN.x1  -0.4181     0.1386  -3.016 0.002837 ** 
---
Signif. codes:  0 '***' 0.001 '**' 0.01 '*' 0.05 '.' 0.1 ' ' 1

Residual standard error: 1.067 on 237 degrees of freedom
Multiple R-squared:  0.09897,   Adjusted R-squared:  0.09136 
F-statistic: 13.02 on 2 and 237 DF,  p-value: 4.333e-06
```

##### 5.1.4 View plot of recapture status and sex over early hibernation fungal loads (Supp Figure 1B)

##### 5.1.5 ***Result:***

- Fungal growth rates over winter are similar between males and
  females that survive WNS.
- Bats that survive winter begin hibernation with lower fungal loads
  than bats that do not survive.

#### 5.2 Do differences in early hibernation roosting temperature explain female biased infections? (Supp Fig 2A)

##### 5.2.1 Are there sex differences in roosting temperature among species?

```
Linear mixed model fit by REML ['lmerMod']
Formula: temp ~ species * sex + (1 | site)
   Data: easpp.t

REML criterion at convergence: 2021.8

Scaled residuals: 
    Min      1Q  Median      3Q     Max 
-5.1106 -0.4507  0.0025  0.5422  3.3058 

Random effects:
 Groups   Name        Variance Std.Dev.
 site     (Intercept) 3.759    1.939   
 Residual             1.075    1.037   
Number of obs: 660, groups:  site, 25

Fixed effects:
                 Estimate Std. Error t value
(Intercept)        8.8851     0.4083  21.762
speciesMYSE        0.1049     0.3681   0.285
speciesPESU        0.5492     0.1909   2.876
speciesEPFU       -1.0462     0.1940  -5.391
sexM               0.1343     0.1209   1.111
speciesMYSE:sexM  -0.4989     0.4461  -1.118
speciesPESU:sexM  -0.1744     0.2295  -0.760
speciesEPFU:sexM  -0.8367     0.2534  -3.302

Correlation of Fixed Effects:
            (Intr) spMYSE spPESU spEPFU sexM   sMYSE: sPESU:
speciesMYSE -0.070                                          
speciesPESU -0.180  0.180                                   
speciesEPFU -0.164  0.147  0.329                            
sexM        -0.211  0.234  0.486  0.469                     
spcsMYSE:sM  0.058 -0.783 -0.129 -0.140 -0.270              
spcsPESU:sM  0.119 -0.118 -0.707 -0.245 -0.531  0.146       
spcsEPFU:sM  0.096 -0.104 -0.209 -0.657 -0.472  0.124  0.245
```

###### 5.2.1.1 View model predicted means (emmean) and standard errors (SE) by sex and species

```
|sex |species |   emmean|        SE|       df|  min.err|  max.err|
|:---|:-------|--------:|---------:|--------:|--------:|--------:|
|F   |MYLU    | 8.885098| 0.4083993| 26.98882| 8.476699| 9.293498|
|M   |MYLU    | 9.019373| 0.4007730| 25.03239| 8.618600| 9.420146|
|F   |MYSE    | 8.990037| 0.5304503| 75.03463| 8.459587| 9.520488|
|M   |MYSE    | 8.625442| 0.4793244| 50.75958| 8.146117| 9.104766|
|F   |PESU    | 9.434322| 0.4184958| 29.78214| 9.015826| 9.852818|
|M   |PESU    | 9.394198| 0.4184027| 29.73164| 8.975795| 9.812600|
|F   |EPFU    | 7.838914| 0.4224191| 30.84960| 7.416495| 8.261333|
|M   |EPFU    | 7.136512| 0.4298539| 33.24702| 6.706658| 7.566365|
```

###### 5.2.1.2 ***Result:***

- No clear difference in early hibernation roosting temperature for
  all species, except EPFU.
- Big brown bat (EPFU) males use cooler temperatures than females,
  however, this species generally uses cooler temperatures and has much
  lower prevalence.

##### 5.2.2 Explore whether addition of temperature term in prevalence model improves predictions.

###### 5.2.2.1 Early hibernation **prevalence** model comparison to models with temperature. All models have site as random effect.

```
|   |Modnames                  |  K|      AIC|  Delta_AIC|  ModelLik|     AICWt|        LL|    Cum.Wt|
|:--|:-------------------------|--:|--------:|----------:|---------:|---------:|---------:|---------:|
|2  |species+sex               |  6| 560.5120|  0.0000000| 1.0000000| 0.4092878| -274.2560| 0.4092878|
|3  |sex * temp+species        |  8| 560.5400|  0.0279976| 0.9860987| 0.4035982| -272.2700| 0.8128860|
|4  |sex+species+temp          |  7| 562.4079|  1.8958481| 0.3875447| 0.1586173| -274.2039| 0.9715033|
|5  |sex * temp+species * temp | 11| 565.8413|  5.3292601| 0.0696251| 0.0284967| -271.9206| 1.0000000|
|1  |null                      |  2| 624.9371| 64.4250934| 0.0000000| 0.0000000| -310.4686| 1.0000000|
```

###### 5.2.2.2 ***Result:***

- Additive model with species and sex is top model (lowest AIC score
  and highest weight), so greater model complexity with a third term is
  not likely warranted.
- Further, in combination with a lack of roosting temperature
  differences between males and females for most species (5.4.1), we are
  unable to identify a consistent effect of temperature on early
  hibernation prevalence that clearly explains sex-specific patterns.
- Therefore, we report the top prevalence model without the
  temperature term.

##### 5.2.3 Now, explore whether addition of temperature term in fungal loads model improves estimates.

###### 5.2.3.1 Early hibernation **fungal load** model comparison with temperature models. All models have site as random effect.

```
|   |Modnames                  |  K|      AIC|  Delta_AIC|  ModelLik|     AICWt|        LL|    Cum.Wt|
|:--|:-------------------------|--:|--------:|----------:|---------:|---------:|---------:|---------:|
|2  |species+sex               |  7| 1434.798|  0.0000000| 1.0000000| 0.4833552| -710.3990| 0.4833552|
|4  |sex+species+temp          |  8| 1435.630|  0.8317078| 0.6597767| 0.3189065| -709.8148| 0.8022616|
|3  |sex * temp+species        |  9| 1437.454|  2.6561511| 0.2649867| 0.1280827| -709.7270| 0.9303443|
|5  |sex * temp+species * temp | 12| 1438.672|  3.8743744| 0.1441087| 0.0696557| -707.3362| 1.0000000|
|1  |null                      |  3| 1484.898| 50.1002131| 0.0000000| 0.0000000| -739.4491| 1.0000000|
```

###### 5.2.3.2 Check whether there is support for temperature term (fl2 = sex+species+temp)

```
Analysis of Deviance Table (Type II Wald chisquare tests)

Response: lgdL
          Chisq Df Pr(>Chisq)    
sex     29.6704  1  5.121e-08 ***
species 45.1656  3  8.532e-10 ***
temp     1.2177  1     0.2698    
---
Signif. codes:  0 '***' 0.001 '**' 0.01 '*' 0.05 '.' 0.1 ' ' 1
```

###### 5.2.3.3 ***Result:***

- Additive model with species and sex as fixed effects is top model
  (lowest AIC score and highest weight), therefore, greater model
  complexity of third temperature term is unlikely to improve model.
- There is no clear support for temperature effect.
- While there may be some additive effect of temperature on fungal
  loads, including it in the model does not improve our ability to explain
  differences beween sexes in early hibernation infection severity.

#### 5.3 Do differences in early hibernation **body mass** explain female biased infections? (Supp Fig 2B)

##### 5.3.1 Early hibernation **prevalence** model comparisons to models with body mass terms. All models have site as a random effect.

```
|   |Modnames                  |  K|      AIC|  Delta_AIC|  ModelLik|     AICWt|        LL|    Cum.Wt|
|:--|:-------------------------|--:|--------:|----------:|---------:|---------:|---------:|---------:|
|2  |species+sex               |  7| 481.8360|   0.000000| 1.0000000| 0.6512378| -233.9180| 0.6512378|
|4  |sex+species+mass          |  8| 483.7974|   1.961359| 0.3750561| 0.2442507| -233.8987| 0.8954885|
|3  |sex * mass+species        |  9| 485.5474|   3.711408| 0.1563429| 0.1018164| -233.7737| 0.9973049|
|5  |sex * mass+species * mass | 13| 492.8109|  10.974899| 0.0041384| 0.0026951| -233.4054| 1.0000000|
|1  |null                      |  2| 624.9371| 143.101131| 0.0000000| 0.0000000| -310.4686| 1.0000000|
```

##### 5.3.2 Early hibernation **fungal loads** model comparisons to models with body mass terms. All models have site as a random effect.

```
|   |Modnames                  |  K|      AIC|  Delta_AIC|  ModelLik|     AICWt|        LL|    Cum.Wt|
|:--|:-------------------------|--:|--------:|----------:|---------:|---------:|---------:|---------:|
|2  |species+sex               |  8| 1270.934|  0.0000000| 1.0000000| 0.4607242| -627.4671| 0.4607242|
|3  |sex+species+mass          |  9| 1271.460|  0.5259779| 0.7687504| 0.3541819| -626.7301| 0.8149061|
|4  |sex * mass+species        | 10| 1272.824|  1.8901462| 0.3886512| 0.1790610| -626.4122| 0.9939671|
|5  |sex * mass+species * mass | 14| 1279.605|  8.6711544| 0.0130943| 0.0060329| -625.8027| 1.0000000|
|1  |null                      |  3| 1309.143| 38.2089652| 0.0000000| 0.0000000| -651.5716| 1.0000000|
```

##### 5.3.3 ***Result:***

- Additive models with sex and species as fixed effects were the top
  models (lowest AIC scores and highest weights) for predicting both
  prevalence and fungal loads compared to models that included body
  mass.
- Adding complexity to the model with a third term of body mass was
  not warranted as it did not improve our ability to explain early
  hibernation infections.

#### 5.4 View plots of early hibernation roosting temperature and mass across species (Supp Fig 2)
