## Appendix; Supplemental Table 2A-2D for "Sex-biased infections scale to population impacts for an emerging wildlife disease": ST2A_est.dat_rev1.htm

| **Supplemental Table 2A. Infection sample summary** | | | | | |
| --- | --- | --- | --- | --- | --- |
| Site | Year | Species | Season | Sex | |
| --- | --- | --- | --- | --- | --- |
|  |  |  |  | Female | Male |
| Illinois | | | | | |
| CKBAL | 2019 | Big brown | Early Hiber |  | 1 |
| CKBAL | 2019 | Little brown | Late Hiber | 1 |  |
| CKBAL | 2019 | Tricolored | Early Hiber | 1 |  |
| CKBAL | 2019 | Tricolored | Late Hiber | 1 |  |
| MERMN | 2016 | Tricolored | Late Hiber | 12 | 1 |
| MERMN | 2018 | Tricolored | Late Hiber | 3 |  |
| MERMN | 2019 | Little brown | Early Hiber |  | 4 |
| MERMN | 2019 | Tricolored | Early Hiber | 3 | 1 |
| MERMN | 2019 | Tricolored | Late Hiber |  | 1 |
| Massachusetts | | | | | |
| R EAT | 2011 | Big brown | Late Hiber | 3 | 7 |
| Michigan | | | | | |
| LOR IN | 2018 | Little brown | Early Hiber | 6 | 16 |
| LOR IN | 2018 | Little brown | Late Hiber | 3 | 2 |
| LOR IN | 2018 | N. long-eared | Early Hiber |  | 1 |
| LOR IN | 2019 | Little brown | Early Hiber | 5 | 16 |
| LOR IN | 2019 | Little brown | Late Hiber | 4 | 15 |
| LOR IN | 2020 | Little brown | Early Hiber | 5 | 18 |
| LOR IN | 2020 | Little brown | Late Hiber | 4 | 17 |
| N ADT | 2018 | Little brown | Early Hiber | 2 | 7 |
| N ADT | 2018 | Little brown | Late Hiber | 4 | 15 |
| N ADT | 2019 | Little brown | Early Hiber | 3 | 16 |
| N ADT | 2019 | Little brown | Late Hiber | 1 | 25 |
| N ADT | 2020 | Big brown | Early Hiber | 8 |  |
| N ADT | 2020 | Big brown | Late Hiber | 9 | 1 |
| N ADT | 2020 | Little brown | Early Hiber | 1 |  |
| N ADT | 2020 | Little brown | Late Hiber | 14 | 30 |
| S C | 2017 | Little brown | Late Hiber | 2 | 12 |
| S C | 2018 | Little brown | Early Hiber | 5 | 8 |
| S C | 2018 | Little brown | Late Hiber | 2 | 1 |
| S C | 2019 | Little brown | Early Hiber | 1 | 10 |
| S C | 2019 | Little brown | Late Hiber | 2 | 10 |
| S C | 2019 | N. long-eared | Early Hiber |  | 1 |
| S C | 2020 | Big brown | Early Hiber | 3 | 1 |
| S C | 2020 | Big brown | Late Hiber | 5 |  |
| S C | 2020 | Little brown | Early Hiber | 2 | 8 |
| S C | 2020 | Little brown | Late Hiber |  | 7 |
| TENSCA | 2018 | Little brown | Early Hiber | 3 | 11 |
| TENSCA | 2018 | Little brown | Late Hiber | 2 | 8 |
| TENSCA | 2018 | N. long-eared | Early Hiber | 1 |  |
| TENSCA | 2019 | Little brown | Early Hiber | 4 | 10 |
| TENSCA | 2019 | Little brown | Late Hiber | 2 | 6 |
| TENSCA | 2020 | Big brown | Early Hiber | 11 | 6 |
| TENSCA | 2020 | Big brown | Late Hiber | 14 | 8 |
| TENSCA | 2020 | Little brown | Early Hiber | 4 | 10 |
| TENSCA | 2020 | Little brown | Late Hiber | 3 | 8 |
| New York | | | | | |
| BROUK | 2011 | Big brown | Late Hiber | 10 | 7 |
| CHCOK | 2011 | Big brown | Late Hiber | 1 | 2 |
| CHCOK | 2011 | Little brown | Late Hiber | 4 | 7 |
| L MIE | 2011 | Big brown | Late Hiber | 5 | 5 |
| L MIE | 2011 | Little brown | Late Hiber | 9 | 6 |
| LIAM H | 2011 | Big brown | Late Hiber | 13 |  |
| LIAM H | 2011 | Little brown | Late Hiber | 8 | 32 |
| LIAM L | 2012 | Big brown | Early Hiber | 8 | 9 |
| LIAM L | 2012 | Big brown | Late Hiber |  | 1 |
| TER IL | 2011 | Big brown | Late Hiber | 11 | 6 |
| TER IL | 2011 | Indiana | Late Hiber |  | 1 |
| TER IL | 2011 | Little brown | Late Hiber | 25 | 28 |
| TER IL | 2012 | Little brown | Late Hiber | 3 | 3 |
| Vermont | | | | | |
| COPER | 2011 | Big brown | Late Hiber | 4 |  |
| Virginia | | | | | |
| ATHIG | 2012 | Big brown | Late Hiber |  | 1 |
| ATHIG | 2012 | Indiana | Early Hiber | 5 | 13 |
| ATHIG | 2012 | Indiana | Late Hiber | 3 | 12 |
| ATHIG | 2012 | Little brown | Early Hiber |  | 1 |
| ATHIG | 2012 | Little brown | Late Hiber | 1 |  |
| ATHIG | 2012 | Tricolored | Early Hiber |  | 1 |
| ATHIG | 2012 | Tricolored | Late Hiber | 3 | 1 |
| BERR | 2012 | Indiana | Early Hiber | 2 | 7 |
| BERR | 2012 | Indiana | Late Hiber | 7 | 20 |
| BERR | 2012 | Little brown | Early Hiber | 6 | 4 |
| BERR | 2012 | Little brown | Late Hiber |  | 8 |
| HERO | 2012 | Big brown | Late Hiber | 2 | 6 |
| HERO | 2012 | Tricolored | Late Hiber | 5 | 2 |
| MANS | 2012 | Big brown | Late Hiber | 3 |  |
| MANS | 2012 | Little brown | Early Hiber | 1 | 10 |
| MANS | 2012 | Little brown | Late Hiber | 10 | 35 |
| MANS | 2012 | Tricolored | Early Hiber | 1 | 2 |
| MANS | 2012 | Tricolored | Late Hiber | 1 | 1 |
| N | 2012 | Indiana | Early Hiber |  | 1 |
| N | 2012 | Little brown | Late Hiber |  | 2 |
| N | 2012 | Tricolored | Early Hiber | 3 | 1 |
| N | 2012 | Tricolored | Late Hiber | 2 |  |
| NELE | 2012 | Little brown | Early Hiber | 1 |  |
| NELE | 2012 | Tricolored | Early Hiber | 1 | 1 |
| NEYS | 2012 | Little brown | Early Hiber | 2 | 4 |
| NEYS | 2012 | Little brown | Late Hiber |  | 1 |
| NEYS | 2012 | Tricolored | Early Hiber | 3 |  |
| NEYS | 2012 | Tricolored | Late Hiber | 5 | 1 |
| RE | 2012 | Tricolored | Late Hiber | 7 | 2 |
| RR CAP | 2012 | Big brown | Late Hiber | 3 |  |
| RR CAP | 2012 | Indiana | Early Hiber | 2 | 7 |
| RR CAP | 2012 | Indiana | Late Hiber | 4 | 10 |
| RR CAP | 2012 | Little brown | Early Hiber | 4 | 10 |
| RR CAP | 2012 | Little brown | Late Hiber | 4 | 14 |
| RR CAP | 2012 | N. long-eared | Early Hiber |  | 1 |
| RR CAP | 2012 | Tricolored | Early Hiber | 3 | 3 |
| RR CAP | 2012 | Tricolored | Late Hiber | 4 | 3 |
| SSRODS | 2012 | Little brown | Early Hiber |  | 1 |
| SSRODS | 2012 | Tricolored | Early Hiber | 4 | 7 |
| SSRODS | 2012 | Tricolored | Late Hiber | 20 | 5 |
| Wisconsin | | | | | |
| CIT | 2019 | N. long-eared | Late Hiber |  | 1 |
| CIT | 2019 | Tricolored | Late Hiber | 2 | 4 |
| CIT | 2020 | Little brown | Early Hiber | 10 | 10 |
| JOH | 2017 | Little brown | Early Hiber | 1 | 4 |
| JOH | 2017 | Little brown | Late Hiber | 2 | 1 |
| JOH | 2018 | Little brown | Late Hiber |  | 1 |
| JOH | 2019 | Little brown | Late Hiber |  | 1 |
| A MIE | 2019 | Tricolored | Early Hiber |  | 1 |
| A MIE | 2020 | Little brown | Early Hiber | 5 | 13 |
| A MIE | 2020 | Tricolored | Early Hiber | 8 | 6 |
| AWBERY | 2017 | Big brown | Late Hiber |  | 1 |
| AWBERY | 2017 | Little brown | Late Hiber | 7 | 5 |
| AWBERY | 2017 | Tricolored | Early Hiber | 2 | 6 |
| AWBERY | 2017 | Tricolored | Late Hiber | 4 | 12 |
| AWBERY | 2018 | Big brown | Late Hiber | 1 | 1 |
| AWBERY | 2018 | Little brown | Late Hiber |  | 1 |
| AWBERY | 2018 | Tricolored | Early Hiber |  | 1 |
| AWBERY | 2018 | Tricolored | Late Hiber | 1 | 2 |
| AWBERY | 2019 | Big brown | Late Hiber | 1 | 4 |
| AWBERY | 2019 | Tricolored | Early Hiber |  | 2 |
| AWBERY | 2020 | Big brown | Early Hiber | 1 | 3 |
| AWBERY | 2020 | Tricolored | Early Hiber | 1 | 1 |
| COBE B | 2018 | Little brown | Early Hiber | 4 | 3 |
| COBE B | 2018 | Little brown | Late Hiber | 4 | 1 |
| COBE B | 2018 | Tricolored | Late Hiber |  | 1 |
| COBE B | 2019 | Little brown | Early Hiber | 2 | 1 |
| COBE B | 2019 | Little brown | Late Hiber | 1 | 2 |
| COBE B | 2019 | N. long-eared | Early Hiber | 1 | 1 |
| COBE B | 2020 | Little brown | Early Hiber | 3 | 1 |
| COBE B | 2020 | Tricolored | Early Hiber |  | 1 |
| DEN OC | 2019 | N. long-eared | Late Hiber | 1 |  |
| DEN OC | 2019 | Tricolored | Late Hiber | 8 | 19 |
| DEN OC | 2020 | Little brown | Early Hiber | 4 | 16 |
| DEN OC | 2020 | Tricolored | Early Hiber | 6 | 10 |
| GER EE | 2018 | Little brown | Early Hiber | 5 | 15 |
| GER EE | 2018 | Little brown | Late Hiber |  | 1 |
| GER EE | 2018 | N. long-eared | Early Hiber | 3 | 9 |
| GER EE | 2018 | Tricolored | Early Hiber | 6 | 9 |
| GER EE | 2019 | Little brown | Late Hiber | 4 | 5 |
| GER EE | 2019 | N. long-eared | Late Hiber |  | 1 |
| GER EE | 2019 | Tricolored | Late Hiber | 4 | 3 |
| GER EE | 2020 | Little brown | Early Hiber | 3 | 1 |
| GER EE | 2020 | N. long-eared | Early Hiber | 1 | 2 |
| GEVIW | 2018 | Little brown | Late Hiber |  | 5 |
| GEVIW | 2018 | Tricolored | Late Hiber | 5 | 2 |
| GEVIW | 2019 | Little brown | Early Hiber |  | 1 |
| GEVIW | 2019 | Little brown | Late Hiber | 1 | 1 |
| GEVIW | 2020 | Little brown | Early Hiber | 2 | 1 |
| IBEL | 2017 | Little brown | Early Hiber | 3 | 7 |
| IBEL | 2017 | Tricolored | Early Hiber | 4 | 6 |
| IBEL | 2018 | Little brown | Early Hiber |  | 4 |
| IBEL | 2018 | Little brown | Late Hiber | 4 | 7 |
| IBEL | 2019 | Little brown | Early Hiber | 4 | 17 |
| IBEL | 2019 | Little brown | Late Hiber | 2 | 1 |
| IBEL | 2019 | N. long-eared | Late Hiber | 1 |  |
| IBEL | 2020 | Little brown | Early Hiber | 4 | 18 |
| LS CLV | 2019 | Big brown | Early Hiber |  | 1 |
| MPFL | 2017 | Tricolored | Early Hiber | 1 | 1 |
| NSTO P | 2018 | Big brown | Early Hiber | 2 |  |
| NSTO P | 2018 | Big brown | Late Hiber | 5 | 1 |
| NSTO P | 2019 | Big brown | Early Hiber | 2 |  |
| NSTO P | 2019 | Big brown | Late Hiber | 1 |  |
| OGRAH | 2019 | Big brown | Early Hiber | 2 | 1 |
| OGRAH | 2019 | Big brown | Late Hiber | 1 |  |
| OGRAH | 2019 | Tricolored | Early Hiber |  | 1 |
| OGRAH | 2020 | Big brown | Early Hiber | 3 | 1 |
| OGRAH | 2020 | Tricolored | Early Hiber |  | 1 |
| OY SAR | 2018 | Big brown | Early Hiber | 1 | 19 |
| OY SAR | 2018 | Big brown | Late Hiber |  | 3 |
| OY SAR | 2018 | Little brown | Late Hiber | 1 | 1 |
| OY SAR | 2019 | Big brown | Early Hiber | 5 | 2 |
| OY SAR | 2019 | Big brown | Late Hiber | 1 |  |
| OY SAR | 2019 | Little brown | Early Hiber | 1 | 18 |
| OY SAR | 2019 | Little brown | Late Hiber | 2 | 3 |
| OY SAR | 2019 | Tricolored | Early Hiber | 6 | 3 |
| OY SAR | 2019 | Tricolored | Late Hiber | 1 |  |
| OY SAR | 2020 | Big brown | Early Hiber | 10 | 6 |
| OY SAR | 2020 | Little brown | Early Hiber | 6 | 14 |
| OY SAR | 2020 | N. long-eared | Early Hiber | 1 |  |
| OY SAR | 2020 | Tricolored | Early Hiber | 2 |  |
| P STTI | 2018 | Tricolored | Early Hiber | 1 |  |
| P STTI | 2018 | Tricolored | Late Hiber | 3 |  |
| P STTI | 2019 | Little brown | Early Hiber | 2 |  |
| R CREK | 2018 | Little brown | Late Hiber | 1 | 2 |
| R CREK | 2018 | Tricolored | Late Hiber | 6 | 2 |
| R CREK | 2019 | Big brown | Late Hiber | 6 |  |
| R CREK | 2019 | Little brown | Late Hiber |  | 1 |
| R CREK | 2019 | Tricolored | Early Hiber | 3 | 4 |
| R CREK | 2019 | Tricolored | Late Hiber | 5 | 1 |
| R CREK | 2020 | Big brown | Early Hiber | 3 |  |
| R CREK | 2020 | Tricolored | Early Hiber | 2 |  |
| SESHE | 2018 | Little brown | Early Hiber | 1 | 1 |
| SESHE | 2018 | Little brown | Late Hiber | 4 | 4 |
| SESHE | 2019 | Big brown | Early Hiber | 1 | 1 |
| SESHE | 2019 | Big brown | Late Hiber | 2 |  |
| SESHE | 2019 | Little brown | Early Hiber | 2 |  |
| SESHE | 2019 | Little brown | Late Hiber | 5 | 12 |
| SESHE | 2019 | Tricolored | Early Hiber |  | 1 |
| SESHE | 2020 | Big brown | Early Hiber | 3 | 1 |
| SESHE | 2020 | Little brown | Early Hiber |  | 9 |
| T RIER | 2017 | Little brown | Early Hiber |  | 1 |
| T RIER | 2017 | Tricolored | Early Hiber | 7 | 2 |
| TH PRT | 2018 | Little brown | Early Hiber | 4 | 28 |
| TH PRT | 2018 | Little brown | Late Hiber | 2 | 4 |
| TH PRT | 2018 | N. long-eared | Early Hiber | 2 | 3 |
| TH PRT | 2018 | N. long-eared | Late Hiber | 1 |  |
| TH PRT | 2018 | Tricolored | Early Hiber | 6 | 12 |
| TH PRT | 2018 | Tricolored | Late Hiber | 2 | 3 |
| TH PRT | 2019 | Big brown | Late Hiber | 1 |  |
| TH PRT | 2019 | Little brown | Early Hiber | 2 | 6 |
| TH PRT | 2019 | Little brown | Late Hiber | 5 | 6 |
| TH PRT | 2019 | Tricolored | Early Hiber | 4 |  |
| TH PRT | 2019 | Tricolored | Late Hiber | 1 |  |
| TH PRT | 2020 | Little brown | Early Hiber |  | 2 |
| TH PRT | 2020 | Tricolored | Early Hiber |  | 1 |
