## Appendix; Supplemental Table 2A-2D for "Sex-biased infections scale to population impacts for an emerging wildlife disease": ST2B_bands_rev1.htm

| **Supplemental Table 2B. Banded bat sample summary** | | | | |
| --- | --- | --- | --- | --- |
| Site | Year | Sex | Nonrecap | Recap |
| --- | --- | --- | --- | --- |
| Michigan | | | | |
| LOR IN | 2016 | Male | 1 | 1 |
| LOR IN | 2017 | Male | 1 | 2 |
| LOR IN | 2018 | Female | 1 | 5 |
| LOR IN | 2018 | Male | 1 | 16 |
| LOR IN | 2019 | Female | 3 | 2 |
| LOR IN | 2019 | Male | 3 | 12 |
| N ADT | 2017 | Male | 1 |  |
| N ADT | 2018 | Female | 2 |  |
| N ADT | 2018 | Male | 2 | 5 |
| N ADT | 2019 | Female | 2 | 1 |
| N ADT | 2019 | Male | 3 | 12 |
| S C | 2018 | Female | 3 | 3 |
| S C | 2018 | Male | 2 | 9 |
| S C | 2019 | Female |  | 1 |
| S C | 2019 | Male | 2 | 9 |
| TENSCA | 2016 | Male | 1 |  |
| TENSCA | 2018 | Female | 2 | 1 |
| TENSCA | 2018 | Male | 3 | 7 |
| TENSCA | 2019 | Female | 2 | 1 |
| TENSCA | 2019 | Male | 4 | 5 |
| Wisconsin | | | | |
| JOH | 2017 | Female | 1 |  |
| JOH | 2017 | Male | 3 |  |
| COBE B | 2018 | Female | 2 | 2 |
| COBE B | 2018 | Male | 2 | 1 |
| COBE B | 2019 | Female | 2 |  |
| COBE B | 2019 | Male |  | 1 |
| GER EE | 2018 | Female | 2 | 2 |
| GER EE | 2018 | Male | 1 | 3 |
| GEVIW | 2019 | Male | 1 |  |
| IBEL | 2018 | Male | 4 |  |
| IBEL | 2019 | Female | 4 | 1 |
| IBEL | 2019 | Male | 19 |  |
| OY SAR | 2019 | Female |  | 1 |
| OY SAR | 2019 | Male | 11 | 3 |
| P STTI | 2019 | Female | 2 |  |
| SESHE | 2018 | Female |  | 1 |
| SESHE | 2018 | Male | 1 |  |
| SESHE | 2019 | Female | 1 | 1 |
| TH PRT | 2018 | Female | 4 |  |
| TH PRT | 2018 | Male | 22 | 3 |
| TH PRT | 2019 | Female | 2 |  |
| TH PRT | 2019 | Male | 4 | 2 |
