## Appendix; Supplemental Table 2A-2D for "Sex-biased infections scale to population impacts for an emerging wildlife disease": ST2C_late_rev1.htm

| **Supplemental Table 3. Population impacts sample summary** | | |
| --- | --- | --- |
| Site | Year | Sample\_size |
| --- | --- | --- |
| Michigan | | |
| LOR IN | 2018 | 5 |
| LOR IN | 2019 | 19 |
| LOR IN | 2020 | 21 |
| N ADT | 2018 | 19 |
| N ADT | 2019 | 26 |
| S C | 2017 | 14 |
| S C | 2018 | 3 |
| S C | 2019 | 12 |
| TENSCA | 2018 | 10 |
| TENSCA | 2019 | 8 |
| Wisconsin | | |
| JOH | 2017 | 3 |
| JOH | 2018 | 1 |
| JOH | 2019 | 1 |
| AWBERY | 2017 | 12 |
| AWBERY | 2018 | 1 |
| COBE B | 2018 | 5 |
| COBE B | 2019 | 3 |
| GER EE | 2018 | 1 |
| GER EE | 2019 | 9 |
| GEVIW | 2018 | 5 |
| GEVIW | 2019 | 2 |
| IBEL | 2018 | 11 |
| IBEL | 2019 | 3 |
| OY SAR | 2018 | 2 |
| OY SAR | 2019 | 5 |
| R CREK | 2018 | 3 |
| R CREK | 2019 | 1 |
| SESHE | 2018 | 8 |
| SESHE | 2019 | 17 |
| TH PRT | 2018 | 6 |
| TH PRT | 2019 | 11 |
