## Appendix; Supplemental Table 2A-2D for "Sex-biased infections scale to population impacts for an emerging wildlife disease": ST2D_fall_rev1.htm

| **Supplemental Table 2D. Fall activity sample summary** | | | |
| --- | --- | --- | --- |
| Site | Year | Females | Males |
| --- | --- | --- | --- |
| CIT | 2020 | 51 | 105 |
| A MIE | 2020 | 7 | 50 |
| A MIE | 2021 | 3 | 38 |
| DEN OC | 2020 | 13 | 36 |
| DEN OC | 2021 | 10 | 22 |
| No detections |  | 46 | 44 |
